# Computational Models Suggest that Human Memory Judgments Exhibit Interference due to the Use of Distributed Representations

**DOI:** 10.1101/2023.04.14.536981

**Authors:** Derek J. Huffman, Ruijia Guan

## Abstract

Episodic memory is a core function that allows us to remember the events of our lives. Given that many events in our life contain overlapping elements (e.g., similar people and places), it is critical to understand how well we can remember the specific events of our lives vs. how susceptible we are to interference between similar memories. Several prominent theories converged on the notion that pattern separation in the hippocampus causes it to play a greater role in processes such as recollection, associative memory, and memory for specific details, while distributed representations in the neocortex cause it to play a stronger role in domain-specific memory. We propose that studying memory performance on tasks with targets and similar lures provides a critical testbed for comparing the extent to which human memory is driven by hippocampal pattern separation vs. more distributed representations (e.g., neocortex) vs. a blend thereof. We generated predictions from several computational models and then tested these predictions in a large sample of human participants. We found a linear relationship between memory performance and target-lure pattern similarity within a neural network simulation of area IT, an object-processing region of the brain. We also observed strong effects of test format on performance and consistent relationships between test formats. Altogether, our results were better accounted for by distributed memory models than the pattern-separated representations of the hippocampus; therefore, our results provide important insight into prominent memory theories by suggesting that memory performance is primarily driven by distributed representations (e.g., neocortex).

---

Episodic memory is a core function that supports and enriches our daily lives. The importance of memory can perhaps be best appreciated by observing patients with Alzheimer’s disease, who experience a devastating loss of self. Therefore, decades of interdisciplinary research have focused on studying the neural basis of memory using a variety of techniques, including direct measures of the brain (e.g., lesion studies, neurophysiology) and computational modeling. An interesting set of competing ideas emerged from these lines of research, with one group of prominent theories suggesting that the hippocampus and medial temporal lobe cortex play highly specialized roles in episodic memory (e.g., recall vs. familiarity in dual-process models; Aggleton & Brown, 1999; Yonelinas, 2002; Eichenbaum, Yonelinas, & Ranganath, 2007) and other theories suggesting that we primarily make memory judgments using a combined or unified memory signal (e.g., Squire, Wixted, & Clark, 2007; Wixted, 2007), which could be implemented by distributed representations (e.g., a rich literature from mathematical psychology; e.g., Murdock, 1982, 1995; Rumelhart, McClelland, & AU, 1986; McClelland, 1986; Hintzman, 1988; Davis, Xue, Love, Preston, & Poldrack, 2014). More generally, a lively debate emerged in the neurosciences about the degree to which memory and perception rely on local representations (e.g., “grandmother cells”, localization of function; cf. Quiroga, Reddy, Kreiman, Koch, & Fried, 2005; Kanwisher, McDermott, & Chun, 1997; Epstein & Kanwisher, 1998) vs. more distributed representations (e.g., Haxby, 2001). The Complementary Learning Systems (CLS) model attempts to unite and integrate these competing frameworks by theorizing that the brain evolved separate neural systems to solve the competing goals of avoiding interference, which is supported by local, pattern-separated representations in the hippocampus, while also allowing for generalization, which is supported by distributed representations in the neocortex (note: the CLS framework is still a dual process model; McClelland, McNaughton, & O’Reilly, 1995; O’Reilly & Norman, 2002; Norman & O’Reilly, 2003; Norman, 2010).

Previous studies have modeled the effects of hippocampal damage on memory performance using the CLS framework, which employs separate neural models of the hippocampus and the neocortex (e.g., medial temporal lobe cortex, which is comprised of areas including the entorhinal cortex, perirhinal cortex, and parahippocampal cortex; Norman & O’Reilly, 2003; Norman, 2010). The CLS model predicts that the hippocampus uses sparse, pattern-separated representations, which allows it to avoid interference, even between memories for similar experiences. In contrast, based on a rich literature in processes such as semantic memory (e.g., Rumelhart & Todd, 1993; McClelland & Rogers, 2003), the CLS model predicts that the neocortex exhibits distributed representations, such that similar experiences cause similar patterns of neural activity, thus allowing organisms to generalize or make predictions about similar experiences. Recognition memory paradigms generally consist of two types of test formats: 1) old/new: a participant sees a single stimulus at a time and decides whether it is old or new (sometimes these formats additionally include a “similar” response), 2) forced-choice: a participant sees two or more images and is asked to select the one that they have seen in the context of the experiment (see Figure 1). The CLS model predicts that hippocampal damage should specifically impair an organism’s ability to discriminate between previously viewed stimuli (i.e., targets) and similar lures but they should exhibit typical performance in discriminating between targets and novel foils during the old/new test format. Specifically, the CLS model predicts that the pattern separation mechanisms of the hippocampus allow it to avoid interference between targets and similar lures whereas the distributed representations in the neocortex exhibit interference between targets and similar lures but can support performance for targets and novel foils (i.e., given the low level of similarity between targets and novel foils). Patients with damage to the hippocampus (Holdstock et al., 2002; Bayley, Wixted, Hopkins, & Squire, 2008; Jeneson, Kirwan, Hopkins, Wixted, & Squire, 2010; Brock Kirwan et al., 2012), and the dentate gyrus in particular (Baker et al., 2016), have been shown to exhibit impaired performance on old/new and old/similar/new tests of targets vs. similar lures but intact performance for targets vs. novel foils, thus supporting the predictions of the CLS model.

**Figure 1:**
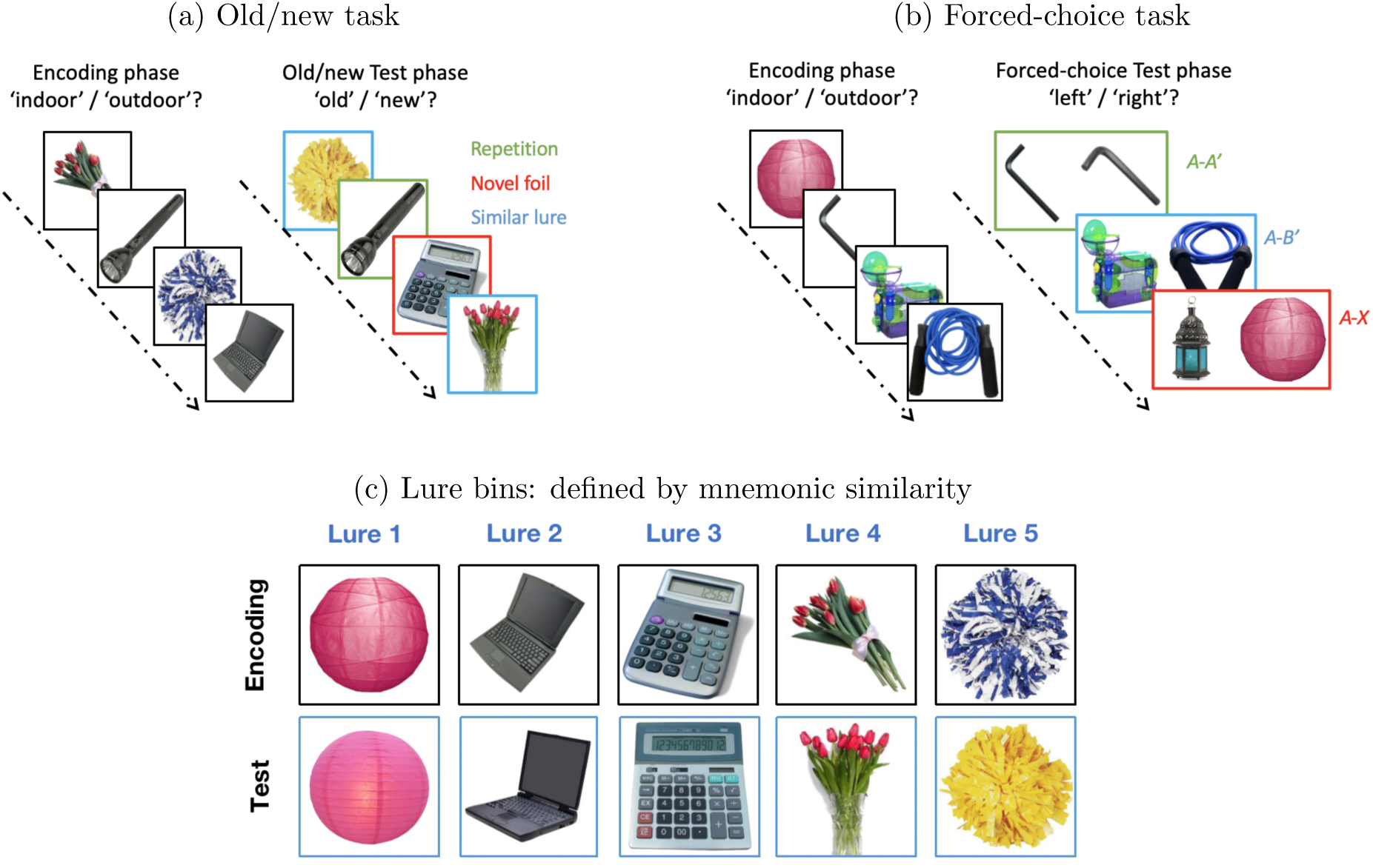
We used two versions of the task and each version contained similar lures that varied as a function of mnemonic similarity. (a) The old/new task was similar the standard versions of the mnemonic similarity task: participants indicated whether each item was “old” or “new”. Participants were instructed to respond “old” only to repetitions and “new” to similar lures and novel foils. (b) We also used a forced-choice version of the mnemonic similarity task with three test formats: A-X (a target and a novel foil), A-A’ (a target and a corresponding similar lure), and A-B’ (a target and a noncorresponding similar lure). (c) In both the old/new and forced-choice tasks, the similar lures varied as a function of mnemonic similarity, which was previously defined as the proportion of times that participants respond “old” to the similar lure

Several neuroscientific studies have supported the prediction that the hippocampus uses pattern separation to reduce the similarity between similar experiences. Foundational studies discovered that the rodent hippocampus contains neurons that respond when the animal occupies a particular location in the environment, termed “place cells” (e.g., O’Keefe & Dostrovsky, 1971; O’Keefe & Nadel, 1978). More recent studies measured the similarity of patterns of activity of hippocampal place cells following changes in the similarity of the spatial environment (e.g., linearly warping from a square to a circular environment; J. K. Leutgeb, Leutgeb, Moser, & Moser, 2007). The theory of hippocampal pattern separation predicts it should exhibit a nonlinear transfer function between the similarity of the inputs (e.g., from high-level cortical areas or the physical/featural similarity) such that it exhibits less similar patterns of activity than its input regions. These studies have generally found that the hippocampus, and the dentate gyrus in particular, exhibits a nonlinear mapping between environmental similarity and place-cell similarity, thus suggesting that the hippocampus is involved in pattern separation (e.g., J. K. Leutgeb et al., 2007; Neunuebel & Knierim, 2014; for review see S. Leutgeb & Leutgeb, 2007).

Studies in humans have further supported the CLS model’s prediction that the hippocampus plays a role in pattern separation. For example, in healthy young adults, functional magnetic resonance imaging (fMRI) has consistently shown the involvement of the hippocampus, and the dentate gyrus and CA3 subregions in particular, in distinguishing between targets (i.e., exact repetitions) and similar lures (i.e., items that are similar to a previously viewed item; e.g., Bakker, Kirwan, Miller, & Stark, 2008; Lacy, Yassa, Stark, Muftuler, & Stark, 2011; Yassa & Stark, 2011; Motley & Kirwan, 2012; Reagh & Yassa, 2014a; Berron et al., 2016). Moreover, these studies have provided evidence for a nonlinear transfer function between DG/CA3 responses and target-lure similarity (e.g., Lacy et al., 2011; Yassa & Stark, 2011; Reagh et al., 2018). Similar to the findings with patients with damage to the hippocampus, healthy aging has been linked to specific disruptions within the hippocampal circuit, including changes in activity in the DG/CA3 subregion of the hippocampus (e.g., hyperactivity: Yassa, Lacy, et al., 2010; Yassa, Mattfeld, Stark, & Stark, 2011; Reagh et al., 2018), disruptions in the integrity of the perforant path (i.e., the input from entorhinal cortex to the hippocampus, as measured by diffusion imaging; Yassa, Muftuler, & Stark, 2010; Bennett & Stark, 2016), and disruptions within the connectivity of a broader hippocampal network (e.g., the integrity of the fornix, a key interface between the hippocampus and neocortex, as measured by diffusion imaging; Bennett, Huffman, & Stark, 2015). Importantly, these hippocampal-network-level disruptions correlated with performance in discriminating between targets and similar lures (on an old/similar/new test format) but did not significantly correlate with performance in discriminating between targets and novel foils, thus further supporting the CLS model’s prediction of a single dissociation in performance for these different types of memory judgments. Moreover, Bakker et al. (2012) and Bakker, Albert, Krauss, Speck, and Gallagher (2015) found that giving a drug to decrease neural excitability, levetiracetam, to patients with amnestic mild cognitive impairment reduced the hyperactivity that was seen in the DG/CA3 subregion in a placebo group of patients (for other studies that have shown hyperactivity in patients with amnestic mild cognitive impairment see: Yassa, Stark, et al., 2010; Tran, Speck, Pisupati, Gallagher, & Bakker, 2017), and the treatment group performed significantly better than the placebo group in discriminating between targets and similar lures (on the old/similar/new test format). Altogether, these findings are consistent with pattern separation in the human hippocampus (for reviews see Yassa & Stark, 2011; Stark, Kirwan, & Stark, 2019).

There is near consensus for the notion that the hippocampus and neocortex represent information differently in the service of memory; however, a critical unaddressed question remains: What is the degree to which we make memory-based decisions based on the pattern-separated representations within the dentate gyrus of the hippocampus vs. the more distributed representations of other hippocampal subregions and the neocortex vs. a composite representation across these systems? While the studies reviewed above suggest that the hippocampus is necessary for typical performance in distinguishing between targets and similar lures and further suggest that the transfer function between targets and similar lures is nonlinear within the dentate gyrus and CA3, other neuroscientific, behavioral, and computational modeling research has suggested that other mechanisms might be used to support memory judgments. For example, in the aforementioned fMRI studies, target-lure similarity was generated via the mnemonic similarity (i.e., memory performance=the proportion of times participants incorrectly responded “old” to similar lures). Therefore, the non-linear mapping between mnemonic similarity and DG/CA3 activity actually suggests a dissociation between memory performance and neural signals. In contrast, activity in CA1 tended to track linearly with mnemonic similarity (e.g., Lacy et al., 2011; Reagh et al., 2018; for review see: Yassa & Stark, 2011), which we think suggests that CA1 exhibits a stronger relationship with behavior. Similarly, immediate early gene studies and single-cell electrophysiology have revealed that the transfer function between the similarity of patterns of active neurons in CA1 and environmental similarity appears to be linear (for review see: Guzowski, Knierim, & Moser, 2004). In fact, evidence from computational models (e.g., Hasselmo, Wyble, & Wallenstein, 1996; Meeter, Murre, & Talamini, 2004) and fMRI (e.g., Kumaran & Maguire, 2006, 2007; Duncan, Ketz, Inati, & Davachi, 2012) has suggested that the hippocampus, and the CA1 subregion in particular, plays a critical role in novelty detection, perhaps through a match-mismatch mechanism. For example, Duncan et al. (2012) used fMRI to show that the human CA1 responded in a linear manner to the degree of changes in an environment. Therefore, these results suggest that the hippocampus might calculate the match vs. mismatch between a past and current experience. Importantly, these kinds of signals would be better instantiated in distributed, overlapping patterns of activity rather than via pattern separation. We will next turn our discussion to distributed memory models and how they have been applied to account for performance on recognition memory tasks.

Different computational models make competing predictions about the nature of the relationship between the similarity of experiences and the resultant memory judgments. The CLS model makes a strong prediction that the hippocampus is generally not useful for computing the global similarity between a probe item and the contents of memory, whereas the neocortex would play a role in calculating the global match (Norman, 2010), which is generally consistent with dual-process models (e.g., Aggleton & Brown, 1999; Yonelinas, 2002; Eichenbaum et al., 2007). Specifically, pattern separation results in a low level of between-item similarity of patterns of activity in the hippocampus if we use lists of items with little overlap between the various items in the stimulus set (i.e., aside from the target-lure similarity, the similarity among the original targets within the list is low), thus the hippocampus largely makes memory judgments based on the similarity between a probe item and one specific target item (i.e., a selective match rather than a global match). However, a class of competing computational models, which we refer to as distributed memory models, suggest that we primarily rely on distributed representations and the global match between a probe item and the contents of our memory to make memory judgments (i.e., similar to the CLS neocortical model; e.g., Murdock, 1982, 1995; Hintzman, 1988). For example, Davis et al. (2014) combined computational modeling and fMRI and they found that recognition memory judgments were correlated with the global similarity between a probe item and the responses to the original target items during encoding within several regions of the medial temporal lobe, including the medial temporal lobe cortex and the hippocampus. Combined with the match-mismatch findings above, these results challenge the notion that the hippocampus cannot compute the global match (but see: LaRocque et al., 2013) and for the hypothesis that pattern separation in the dentate gyrus would result in a direct behavioral readout on memory tasks. Moreover, given the strong relationship between global match and performance (Davis et al., 2014), these results provide more direct evidence that memory judgments might be supported by a global matching mechanism (note: they did not specifically examine similar lures, which is a primary aim of our work here).

Performance on forced-choice recognition memory tasks has often been found to be better supported by predictions of distributed memory models (e.g., related to the neocortex) than by the pattern separation mechanisms of the dentate gyrus. Specifically, there are three basic types of forced-choice test formats: A-X (a target and an unrelated, novel foil; traditional forced-choice recognition), A-A’ (a target and a corresponding similar lure; e.g., a picture of an apple and a similar apple), and A-B’ (a target and a noncorresponding similar lure; e.g., a picture of an apple and a basketball that is similar to one that was studied in the encoding phase; note, we use the nomenclature of Tulving, 1981; see Figure 1). The CLS framework predicts that damage to the hippocampus will cause a test-format-specific impairment in discriminating between targets and similar lures within forced-choice tests: patients should be impaired on the A-B’ test format but they should exhibit typical performance on the A-A’ test format as well as the A-X test format (see simulation 4 from Norman & O’Reilly, 2003; Norman, 2010). In contrast to the findings for the old/new test formats, the support for the predictions of selective impairments on the forced-choice formats has been equivocal. A case study of a single patient suggested that damage to the hippocampus specifically impaired performance in discriminating between targets and similar lures in the old/new test format with intact performance on the A-A’ test format (Holdstock et al., 2002; for similar findings in patients with amnestic mild cognitive impairment see: Westerberg et al., 2006). However, other studies with a larger sample of patients suggest that hippocampal damage impairs performance on both the old/new and the A-A’ test formats (Bayley et al., 2008; Jeneson et al., 2010). Moreover, Jeneson et al. (2010) found that patients with selective hippocampal damage exhibited a similar degree of impairment on the A-A’ test format as the A-B’ test format. Specifically, they did not observe a significant patient-group by test-format interaction and they observed that the patients were significantly impaired on both test formats (as well as the old/new test format). Therefore, these latter results provide evidence that is discordant with the prediction of the standard version of the CLS hippocampal model, and we turn next to discussing performance on the forced-choice test formats in healthy young adults.

The support for the CLS hippocampal model’s prediction of similar performance on the A-A’ and A-B’ test formats in healthy young adults (i.e., given their intact hippocampal circuit; see see Figure 2a; cf. Migo, Montaldi, Norman, Quamme, & Mayes, 2009) has also been equivocal (but note that Norman & O’Reilly, 2003; Norman, 2010 discussed the idea that human memory would likely better be modeled by combining the output of the hippococampal and medial temporal lobe cortex model, which we will return to in the General Discussion). Migo et al. (2009) found that healthy young adults exhibit similar performance on the A-A’ and the A-B’ forced-choice test formats (i.e., the Recognition condition from Experiment 1 in Migo et al., 2009). However, other, larger studies have shown clear effects of test format (e.g., Experiment 1 and 2 from Huffman & Stark, 2017; Hintzman, 1988; Tulving, 1981; Rollins, Khuu, & Lodi, 2019; Fandakova, Johnson, & Ghetti, 2021). Specifically, the general finding in these studies is that performance on the A-A’ test format is significantly better than performance on the A-B’ test format, and this finding is consistent within healthy young adults (Experiment 1 and 2 from Huffman & Stark, 2017; Hintzman, 1988; Tulving, 1981; Rollins et al., 2019; Fandakova et al., 2021), healthy middle-aged adults (Jeneson et al., 2010), healthy older adults (Experiment 1 from Huffman & Stark, 2017), and patients with damage to the hippocampus (Jeneson et al., 2010). Altogether, the evidence for the hippocampal model’s prediction of a null effect of a difference between the A-A’ and the A-B’ forced-choice test format (cf. Migo et al., 2009) is tenuous. We suggest that more evidence points toward a significant effect of test format because several published studies have shown a significant A-A’ *>* A-B’ test format effect and it is difficult to interpret null effects in the other papers (especially without a Bayesian analysis). Importantly, distributed memory models readily account for the A-A’ *>* A-B’ test format effect (e.g., Hintzman, 1988; Huffman & Stark, 2017; also see the neocortical model from Simulation 4 in Norman & O’Reilly, 2003).

**Figure 2:**
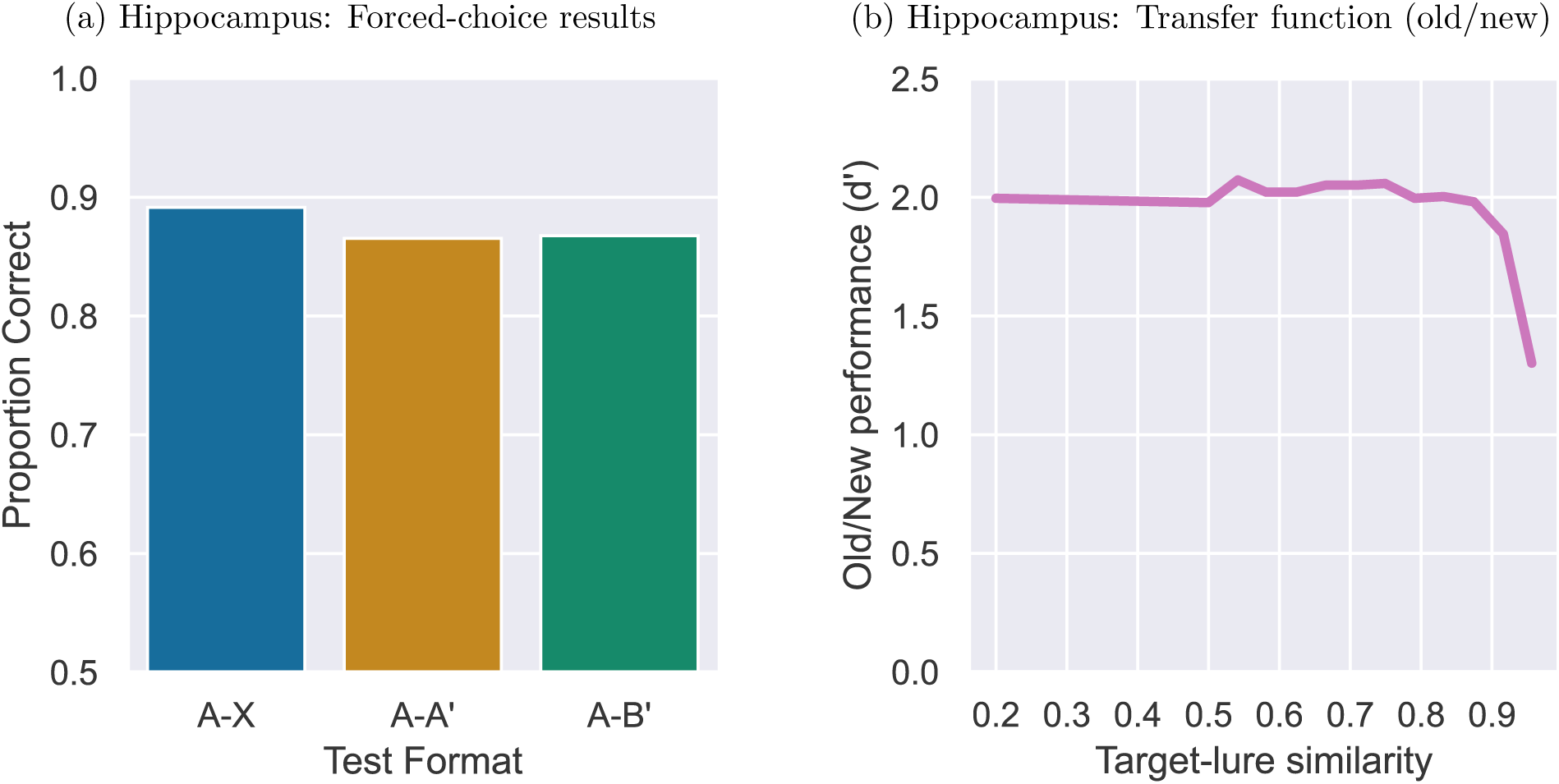
(a) Norman and O’Reilly (2003) found that the simulated behavioral performance of the CLS hippocampal model was not significantly different on the A-A’ vs. A-B’ forced-choice test formats (replotted from the 0.71 overlap condition in Figure 12 of Norman & O’Reilly, 2003), and there were cases in which performance on the A-B’ test format was actually better than the A-A’ test format. (b) Norman and O’Reilly (2003) found that the simulated behavioral performance of the CLS hippocampal model on the old/new test format was highly resilient to target-lure similarity, such that there was virtually no difference in performance for discriminating between targets vs. novel foils (target-lure similarity of 0.2) or targets vs. similar lures with a moderate to high level of overlap (i.e., similarity from 0.5 to 0.875). In fact, the hippocampal model did not exhibit interference between targets vs. similar lures until extremely high levels of similarity (i.e., similarity *>* 0.875; replotted from Figure 9 of Norman & O’Reilly, 2003).

We aimed to test contradictory predictions from distributed memory models and the CLS hippocampal model about performance on memory tasks with targets, similar lures, and novel foils to determine whether memory is driven primarily by pattern separation in the hippocampus vs. distributed representations (e.g., neocortex) vs. a composite representation between these systems. First, distributed memory models and the CLS hippocampal model make contradictory predictions about the relationship between memory performance and the similarity between targets and similar lures. Specifically, as we show here, distributed memory models predict that the degree of similarity will cause a relatively strong, continuous, and monotonic relationship between performance in discriminating between targets and similar lures. In contrast, the CLS hippocampal model predicts a much more modest effect of target/lure similarity, which exhibits a relatively flat or null effect of stimulus similarity until the target/lure similarity is extremely high, given its use of pattern separation in the dentate gyrus and CA3 (see Figure 2b and Norman & O’Reilly, 2003). We aimed to quantify a measure of physical or semantic similarity between our target and lure items using a biologically inspired artificial neural network to simulate patterns of activity in the inferior temporal cortex (area IT; Kubilius et al., 2019), which is a high-level visual region involved in object recognition and it is heavily interconnected with the perirhinal cortex (Suzuki & Amaral, 1994). Thus, we argue that the artificial neural network allows us to empirically determine the transfer function of performance on the task vs. an objective measure of the target/lure similarity. We examined the effects of target/lure similarity on performance across various test formats, including the old/new, A-A’, and A-B’ test formats. Second, we tested for the A-X *>* A-A’ *>* A-B’ test format effect, which should be significant if humans primarily make memory judgments by relying on distributed representations but null if humans primarily make memory judgments using the pattern-separated representations of the hippocampus. We further aimed to test the relationship between performance on the forced-choice and old/new test formats to test specific predictions of distributed memory models. Finally, we aimed to further generate and test predictions from distributed memory models and neural networks to test how they compare with human performance. To foreshadow our findings and conclusions, we provide evidence that performance across several test formats that ask participants to distinguish between targets, similar lures, and novel foils is more consistent with the distributed memory models (vs. hippocampal pattern separation), thus providing important insight into neural and computational theories of human memory and the behavioral tasks that we use to study these systems.

## Quantification of Target/Lure Similarity with an Artificial Neural Network

We used an artificial neural network to quantify the similarity between our target and lure items. Briefly, previous research has focused on defining the similarity between targets and lures using a measure of mnemonic performance on the task (Lacy et al., 2011; Stark, Yassa, Lacy, & Stark, 2013; Stark, Stevenson, Wu, Rutledge, & Stark, 2015; Stark et al., 2019). Specifically, these studies have combined items into lure bins by quantifying the proportion of times that a participant makes an “old” response to a similar lure item, with a higher proportion of “old” responses indicating higher mnemonic similarity. These approaches are very helpful for quantifying and comparing the mnemonic similarity between stimuli. However, we think that it would also be interesting to study the relationship between a measure of stimulus similarity (i.e., a physical or semantic measure of similarity between targets and lures) to compare the transfer function between this objective measure of stimulus similarity and the degree of memory interference for the similar lures. Thus, we modeled pattern similarity using an artificial neural network of area IT, a high level visual area and one of the primary inputs into perirhinal cortex (Suzuki & Amaral, 1994).

### Methods

We extracted features (i.e., modeled neural responses) from the 3rd convolutional layer of IT (i.e., the penultimate layer) from CORnet-S (Kubilius et al., 2019) using the THINGSvision toolbox (Muttenthaler & Hebart, 2021). Briefly, CORnet-S has been shown to provide a good approximation both to neural responses in IT as well as to behavioral data across several studies (Kubilius et al., 2019; Muttenthaler & Hebart, 2021). Moreover, area IT is shown to provide the main inputs into perirhinal cortex (Suzuki & Amaral, 1994), and thus the simulated activity in IT can be seen as an approximation of the input to the medial temporal lobe for object-memory tasks.

THINGSvision automatically resizes images to 256 pixels using bilinear interpolation and then performs a center crop to 224×224 pixels. Therefore, we changed our images to all be a square, while maintaining image proportions, by applying a white border (note all images in the Mnemonic Similarity Task are objects on a white background). Here, we set the white border to be 256 / 224 times the maximum dimension of the image (i.e., to avoid the model cropping part of the image). After extracting the simulated responses from IT, we quantified the similarity between the target-lure pairs by calculating Pearson’s correlation coefficient between the model’s response to the target item and the lure item (i.e., we calculated the pattern similarity; Kriegeskorte, 2008). We then calculated Pearson’s correlation coefficient between the resultant pattern similarity scores and 1-p(“old”) data from previous reports (Stark et al., 2013, 2015, 2019). Here, we were specifically interested in comparing whether the transfer function between mnemonic discrimination and target/lure similarity is monotonic and relatively continuous (i.e., in line with the distributed memory models; MINERVA 2: Figure 5b, TODAM: Figure 6b, the familiarity-based Hopfield/autoassociative neural network: Figure 7b, and the recall-based Hopfield/autoassociative neural network: Figure 8b) or whether it is solely driven by only extremely high similarity values (i.e., target-lure similarity values that are greater than 0.875, which is what the CLS hippocampal model predicted; see Figure 2b; Norman & O’Reilly, 2003). Accordingly, we compared the fit of various powers of the following power function:

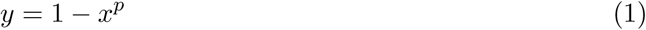

where *x* indicates the pattern similarity from CORnet-S and *p* indicates the power of the function. When *p* is set to 1, the predicted behavioral performance is a linear function based on the theoretical “input” similarity in IT (i.e., more in line with the distributed memory models; see Figure 3). When *p* is set to higher values, the predicted behavioral performance is a nonlinear function, which would theoretically indicate greater pattern separation following the representation in IT (i.e., more in line with the CLS hippocampal model; see Figure 2b and the greater power values in Figure 3). We compared the Bayesian information criteria (BIC) between models with different values of *p* (i.e., because zero parameters differed), by comparing both the raw BIC values (i.e., lower BIC indicated the preferred model) as well as the approximation of Bayes factors using the following equation (Wagenmakers, 2007; Jarosz & Wiley, 2014):

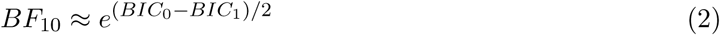

**Figure 3:**
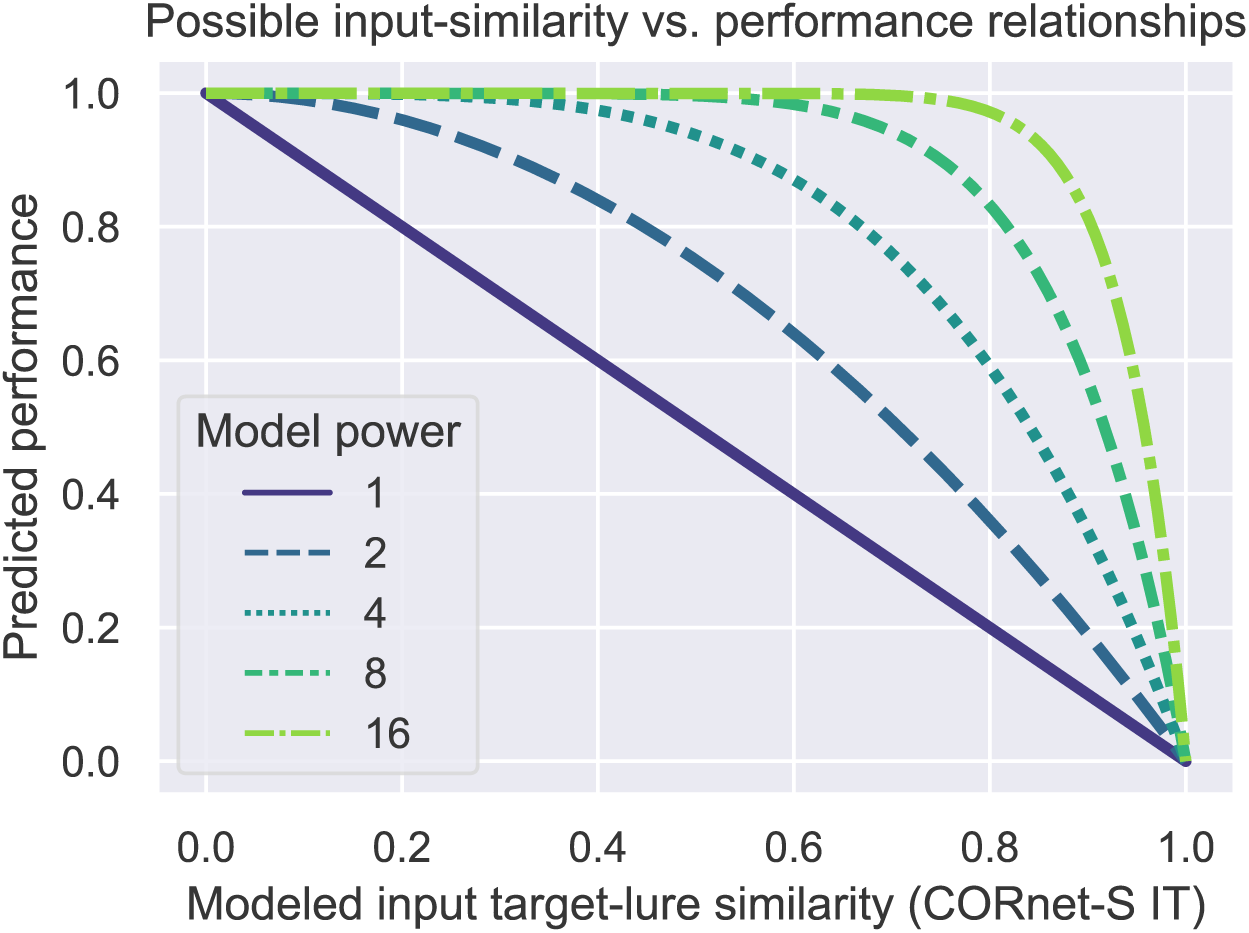
Possible input-similarity vs. performance relationships. Distributed memory models (e.g., based on the neocortex) would predict a monotonic and perhaps relatively linear relationship between the modeled input target-lure similarity (IT within CORnet-S) and performance (i.e., model power values near 1). Conversely, if humans rely on hippocampal pattern separation for performance, then we would expect to see a nonlinear relationship between the modeled input target-lure similarity and performance (i.e., model power values greater than 1 and higher values would reflect greater degrees of pattern separation; e.g., model power values around 16 are similar to the predictions of the CLS hippocampal model, see Figure 2b).

**Figure 4:**
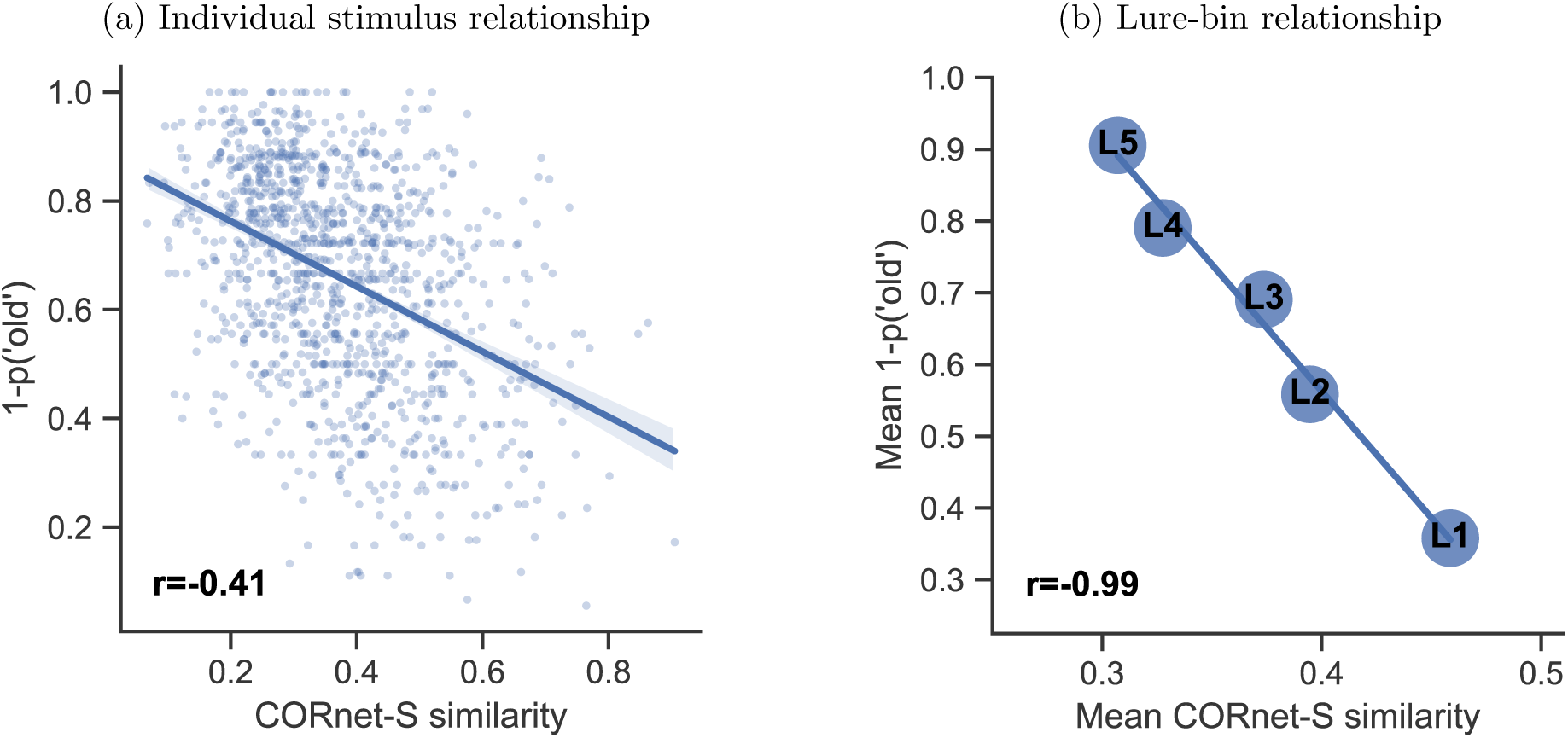
We observed a significant correlation between the simulated target-lure pattern similarity in the IT of CORnet-S and human mnemonic discrimination performance in the data that were used to define the original lure bins (Lacy et al., 2011; Stark et al., 2019, 2013, 2015). (a) We observed a significant correlation between CORnet-S similarity and human memory performance (1-p(‘old’)) for individual stimulus pairs. Here, the correlation between the model and behavior compares favorably to other studies that have compared pattern similarity in artificial neural networks and behavioral performance (e.g., on perceptual tasks). (b) We also averaged the pattern similarity and the human memory performance values (1-p(‘old’)) within each lure bin to reduce noise, and we again found a striking relationship between pattern similarity and performance on the task. The points indicate the lure bin of each data point (see Figure 1). Altogether, these data support the predictions of the distributed memory models (see Figure 5b, Figure 6b, Figure 7b, Figure 8b) but are at odds with the predictions of the CLS hippocampal model (see Figure 2b).

**Figure 5:**
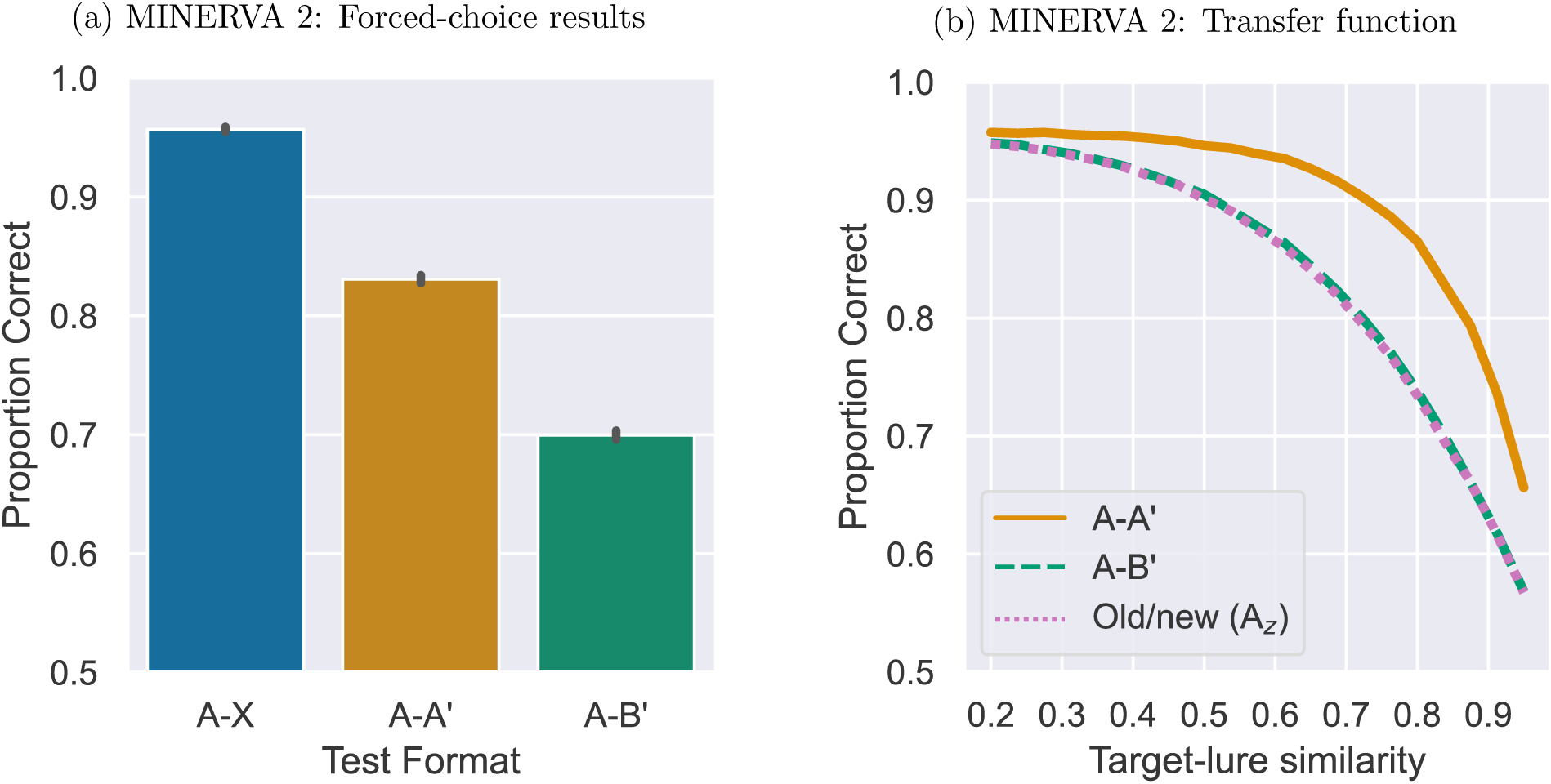
(a) MINERVA 2 predicts a clear test-format effect: A-X *>* A-A’ *>* A-B’ (note the difference from the predictions of the CLS hippocampal model). (b) MINERVA 2 predicts a clear, monotonic relationship between target-lure similarity and performance on the A-A’, A-B’, and old/new (*A_z_*) test formats (note the difference from the predictions of the CLS hippocampal model, which predicts that performance on the old/new test format will only exhibit interference at very high levels of target-lure similarity). Each of these simulations included 1,000 simulated participants. Error bars indicate the 95% confidence intervals. Please compare these results to the empirical results in Figure 4, Figure 12, and Figure 13.

**Figure 6:**
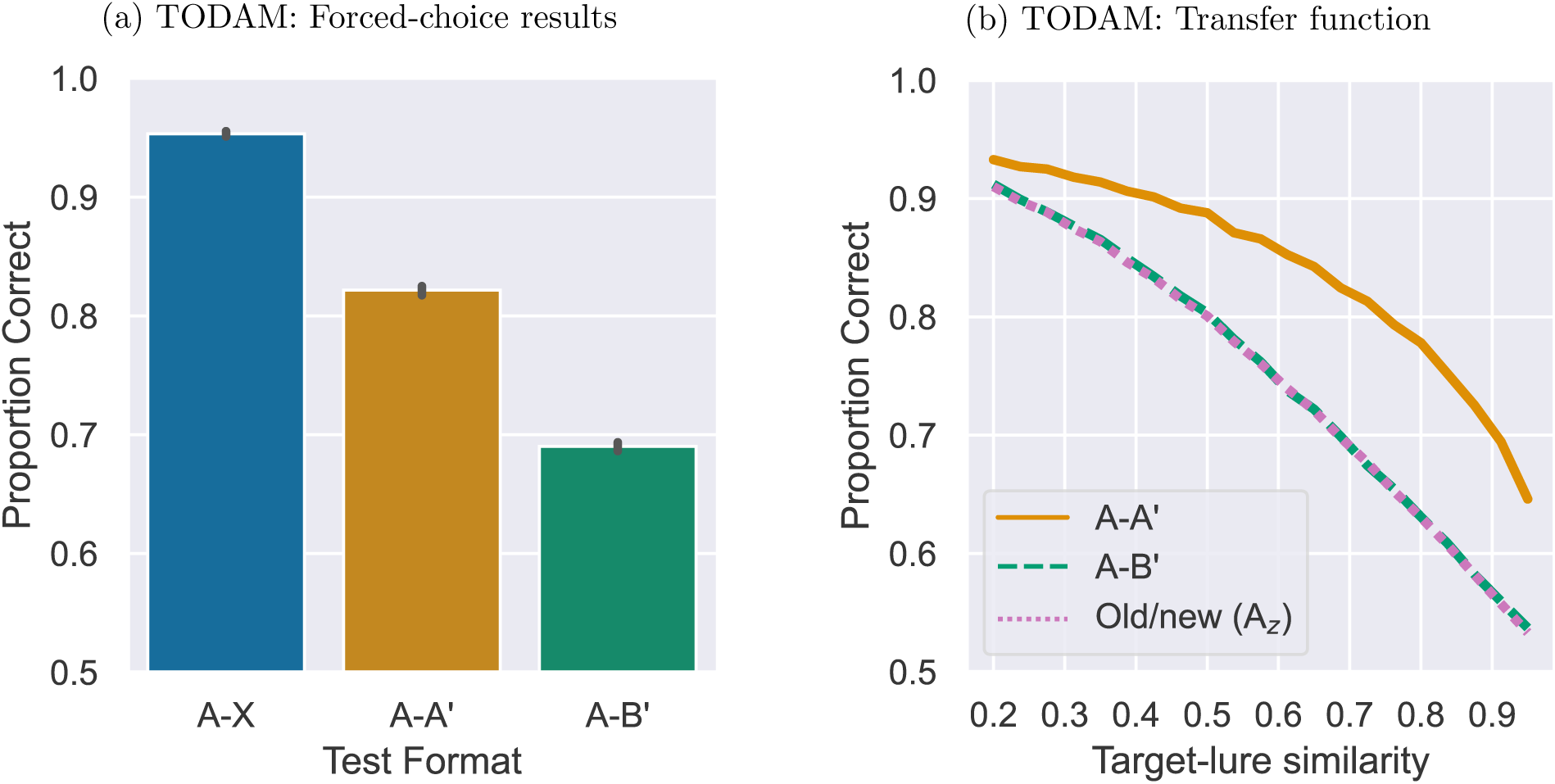
(a) TODAM predicts a clear test-format effect: A-X *>* A-A’ *>* A-B’ (note the difference from the predictions of the CLS hippocampal model). (b) TODAM predicts a clear, monotonic relationship between target-lure similarity and performance on the A-A’, A-B’, and old/new (*A_z_*) test formats (note the difference from the predictions of the CLS hippocampal model, which predicts that performance on the old/new test format will only exhibit interference at very high levels of target-lure similarity). Each of these simulations included 1,000 simulated participants. Error bars indicate the 95% confidence intervals. Please compare these results to the empirical results in Figure 4, Figure 12, and Figure 13.

**Figure 7:**
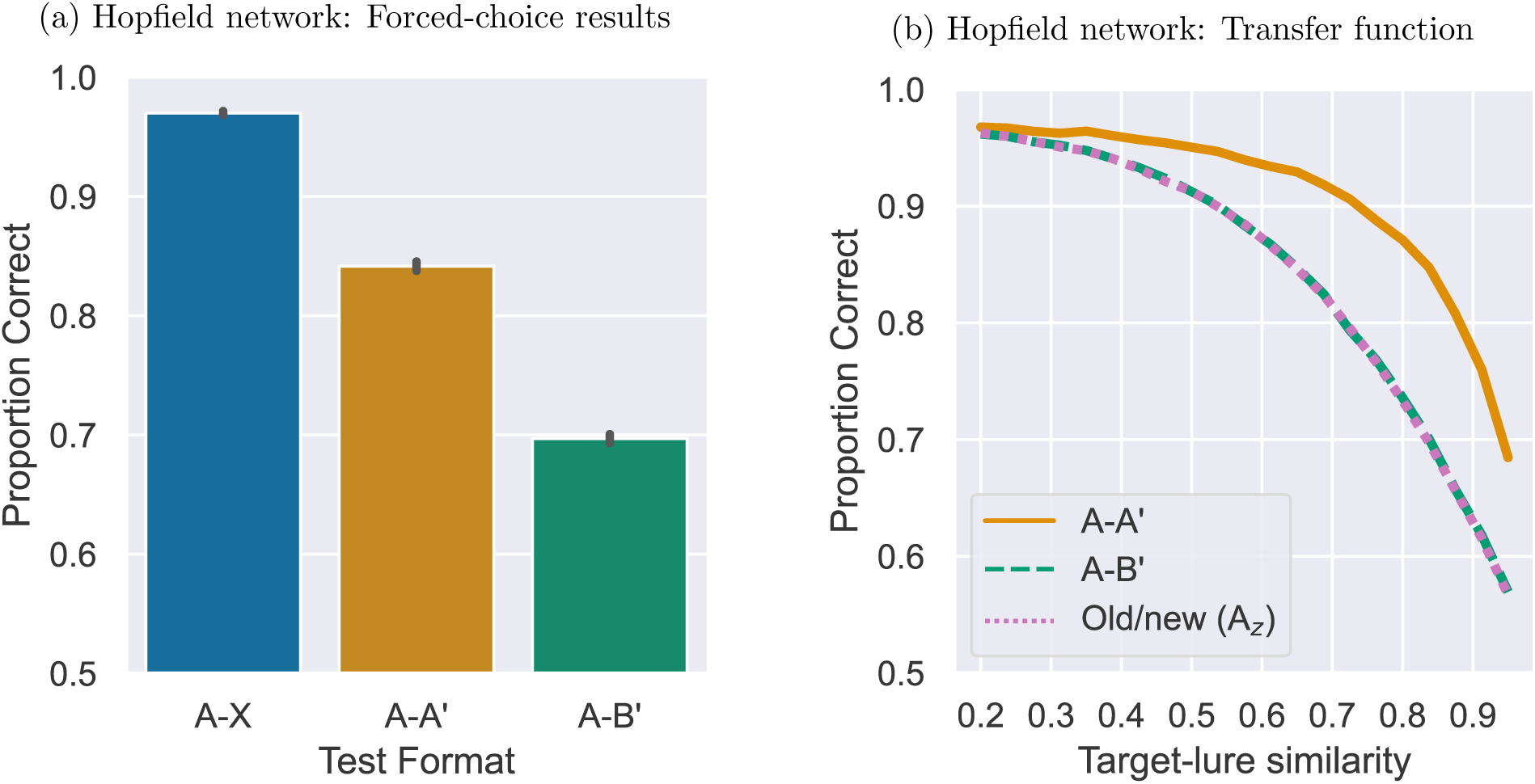
(a) The familiarity-based version of the Hopfield/autoassociative neural network predicts a clear test-format effect: A-X *>* A-A’ *>* A-B’ (note the difference from the predictions of the CLS hippocampal model). (b) The familiarity-based version of the Hopfield/autoassociative neural network predicts a clear, monotonic relationship between target-lure similarity and performance on the A-A’, A-B’, and old/new (*A_z_*) test formats (note the difference from the predictions of the CLS hippocampal model, which predicts that performance on the old/new test format will only exhibit interference at very high levels of target-lure similarity). Each of these simulations included 1,000 simulated participants. Error bars indicate the 95% confidence intervals. Please compare these results to the empirical results in Figure 4, Figure 12, and Figure 13.

**Figure 8:**
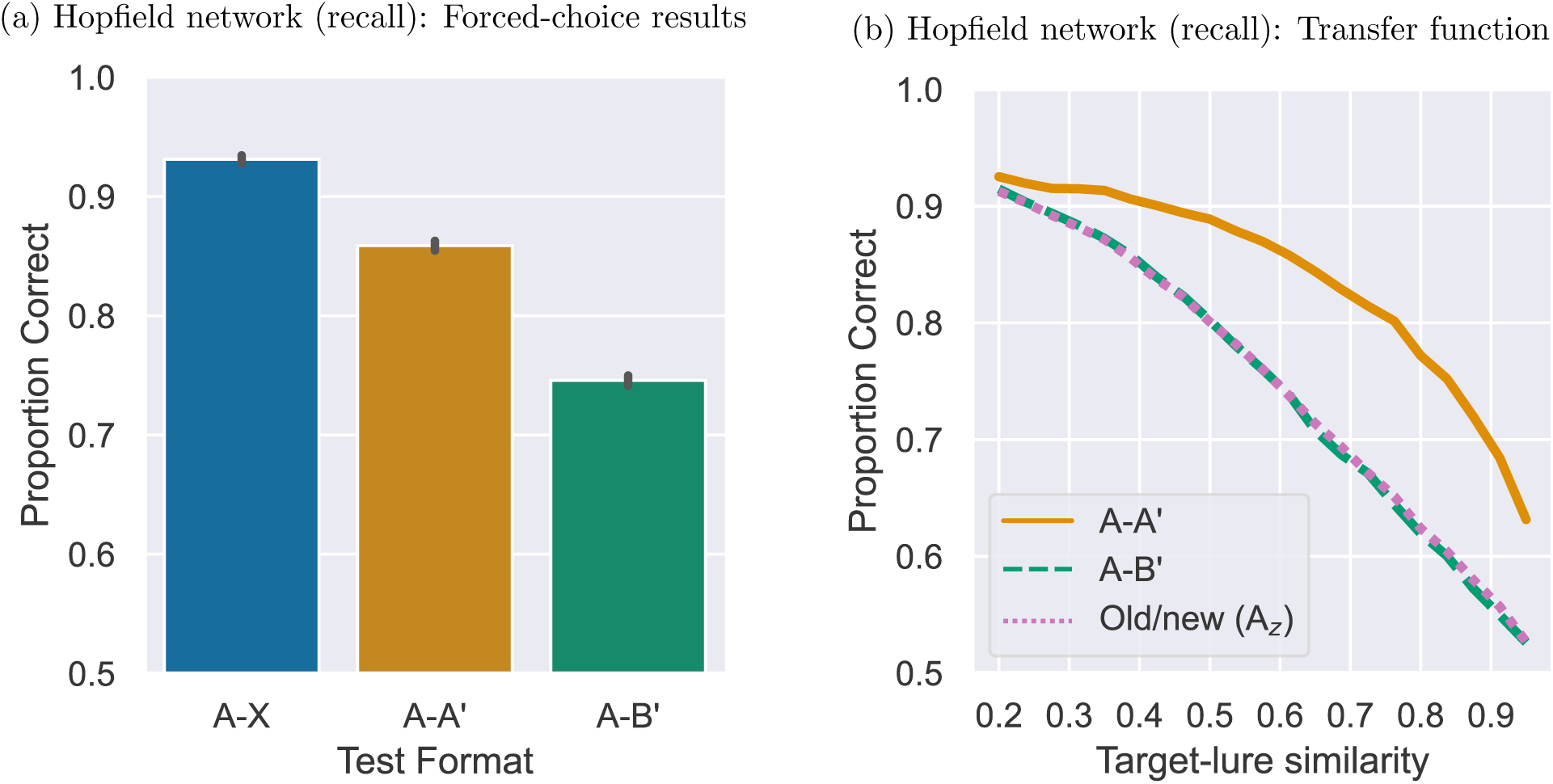
(a) The recall-based version of the Hopfield/autoassociative neural network predicts a clear test-format effect: A-X *>* A-A’ *>* A-B’ (note the difference from the predictions of the CLS hippocampal model). (b) The recall-based version of the Hopfield/autoassociative neural network predicts a clear, monotonic relationship between target-lure similarity and performance on the A-A’, A-B’, and old/new (*A_z_*) test formats (note the difference from the predictions of the CLS hippocampal model, which predicts that performance on the old/new test format will only exhibit interference at very high levels of target-lure similarity). Each of these simulations included 1,000 simulated participants. Error bars indicate the 95% confidence intervals. Please compare these results to the empirical results in Figure 4, Figure 12, and Figure 13.

Here, we compared the fit of *p* values of 1 vs. 2, 4, 8, and 16.

### Results and Discussion

We first aimed to test the nature of the transfer function between human memory performance and target-lure similarity. Specifically, as we discussed in the Introduction, the CLS version of the hippocampal model suggests that the transfer function between performance and the target-lure similarity is highly nonlinear (Norman & O’Reilly, 2003; Norman, 2010). Specifically, this theory suggests that the hippocampus will largely avoid interference until the target-lure similarity is very high (i.e., above 87.5% overlap; see Figure 2b). In contrast, the distributed memory models predict that the target-lure similarity will more strongly and continuously affect performance (see below; also see the cortical model from Norman & O’Reilly, 2003). Therefore, determining the transfer function between performance and target-lure similarity would provide evidence to suggest whether humans primarily based on the pattern-separated representations of the hippocampus vs. distributed representations (e.g., the neocortex) vs. a composite representation across these systems. Here, we used an artificial neural network to simulate responses in area IT to all of the images in 6 stimulus sets of the Mnemonic Similarity Task (Stark et al., 2013, 2015, 2019), which each contained 192 target-lure pairs (i.e., a total of 1152 images).

As we discuss in the Methods, we extracted patterns of activity from the 3rd convolutional layer of IT from CORnet-S (i.e., the penultimate layer; Kubilius et al., 2019; Muttenthaler & Hebart, 2021) and calculated the pattern similarity (Pearson correlation) between the responses to the target and the lure images. We compared these pattern-similarity values to behavioral performance, which we defined here based on the proportion of times that participants made an “old” response to the lures (1-p(‘old’)), as in past research (Stark et al., 2013, 2015, 2019).

We observed a strong linear relationship between performance on the task and the simulated pattern similarity (Pearson’s *r* = −0.41, *p* = 1.71 × 10*^−^*^48^; see Figure 4). Note that the magnitude of our observed effect compares favorably to other approaches linking neural network models with human behavior on perceptual tasks (Kubilius et al., 2019; Muttenthaler & Hebart, 2021). Moreover, we averaged the pattern similarity and performance within each lure bin to reduce noise (i.e., because the simulations are not a perfect recapitulation of actual neural responses), and we again observed a strong linear relationship between these variables (Pearson’s *r* = −0.99). Additionally, we compared the fit of various models of power relationships between the CORnet-S pattern similarity and behavioral performance (see Equation (1) and Figure 3), where greater power values would theoretically reflect greater reliance on pattern separation following the putative “input” of information to the medial temporal lobe from IT. We found that the linear model (*BIC* = −656.18) provided a better fit than models with power values of 2 (*BIC* = −645.34; *BF*_10_ ≈ 226.8), 4 (*BIC* = −584.13; *BF*_10_ ≈ 4.45 × 10^15^), 8 (*BIC* = −493.33; *BF*_10_ ≈ 2.32 × 10^35^), and 16 (*BIC* = −454.01; *BF*_10_ ≈ 8.02 × 10^43^).

Our results suggest that the transfer function between mnemonic discrimination performance and target-lure similarity exhibits a strong, continuous, and perhaps linear relationship. Moreover, while it is difficult to interpret the pattern similarity values directly, the magnitude of these values span a broad range and performance is not solely impaired by only the most similar lures (e.g., pattern similarity greater than 0.75). Altogether, these results suggest that human performance is primarily driven by distributed representations (e.g., neocortex of hippocampal subregions CA1 or CA3; see following sections) rather than the pattern-separated representations of the hippocampus (e.g., the dentate gryus; e.g., Norman & O’Reilly, 2003; Norman, 2010; see Figure 2b) or a hybrid, composite relationship of representations across the combined models.

## Comparison of Predictions from the Computational Models

We next aimed to generate predictions from several computational models regarding performance on memory tasks with targets, similar lures, and novel foils (e.g., the Mnemonic Similarity Task; Kirwan & Stark, 2007; Stark et al., 2013, 2015, 2019; see Figure 1). Here, we specifically compared the predictions from the CLS model of the hippocampus (Norman & O’Reilly, 2003; Norman, 2010) with several models, including MINERVA 2 (a multiple-trace global-matching model; Hintzman, 1984, 1988), theory of distributed and associative memory (TODAM; a unitrace global-matching model; Murdock, 1982, 1995), and two implementations of a Hopfield/autoassociative neural network (similar computational architecture to the CA3 subregion of the hippocampus: Rolls, 2007; Hopfield, 1982; Rizzuto & Kahana, 2001). Given that all of these models implement distributed representations, we will refer to these collectively as “distributed memory models” (which also stands in contrast to the CLS hippocampal model, which emphasizes sparse, pattern-separated representations). We then tested the predictions from these models with empirical performance in a large sample of human participants.

### Computational Modeling Methods

#### Global Matching Models: MINERVA 2 and TODAM

We previously found that MINERVA 2 and TODAM can account for many aspects of performance on the forced-choice versions of the Mnemonic Similarity Task (Huffman & Stark, 2017; cf. Hintzman, 1988). Specifically, these models suggest that performance on the tasks will vary as a function of test format: A-X *>* A-A’ *>* A-B’, similar to our previous empirical data. Here, we again used these models to replicate the finding of the test format effect. Additionally, we extended these models to make novel predictions, which we tested in our empirical data.

We modeled both MINERVA 2 (Hintzman, 1984, 1988) and TODAM (Murdock, 1995) in Python. To model items in MINERVA 2, we drew random integers of -1, 0, and 1 using the randint function from numpy. To generate similar lures, we randomly redrew from the random integers for a specified proportion of the features using the binomial function from numpy to select the features to be changed. We modeled learning by having the model learn each feature with probability of *L* and to not learn each feature with probability 1 - *L*. Specifically, we used the binomial function from numpy to select which features would be remembered for each trial. We then multiplied this binomial matrix by the encoding list to simulate the learning phase. Thus, the learning phase generated a matrix with M rows (i.e., memory traces) and N columns (i.e., features). During the testing phase, we calculated the summed similarity of the probe to the memory matrix using the following equations (Hintzman, 1984, 1988):

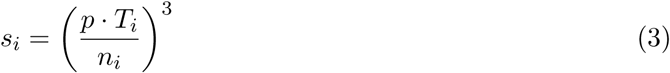

where p is the probe item, *T_i_* is one of the items in memory (i.e., one of the rows of the memory matrix), and *n_i_*is the number of features that are nonzero in either *p* or *T_i_*(i.e., the part of the equation within the parentheses is a normalized dot product, which calculates the match between the probe and a given memory trace and normalizes by the number of nonzero features). The cubing function makes the match nonlinear, which can be thought of as a form of pattern separation. We then summed the similarity values across all items in the memory matrix using the following equation (Hintzman, 1984, 1988):

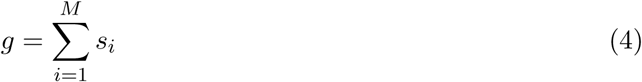

The encoding phase of MINERVA 2 is an instantiation of a multiple-trace model. However, the nature of the way that memory retrieval is modeled makes it a global matching model (i.e., because it sums the similarity across all memory traces to calculate the overall memory match for each probe item).

We modeled TODAM using the item-only version of the model (Murdock, 1995); however, TODAM has traditionally focused on modeling associative memory (e.g., Murdock, 1982). Items in TODAM are represented as vectors, similar to MINERVA 2. Each feature of the item vectors in TODAM is generated as a random draw from a normal distribution with mean 0 and standard deviation 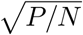, where *N* is the number of features (i.e., the length of the item vector) and in our case *P* is equal to 1. Note that setting the standard deviation with *P* equal to 1 has the desirable effect of making the item vectors have approximately unit length, thus allowing us to calculate the similarity using the dot product (i.e., this has a normalizing effect on the calculation of the dot product; Murdock, 1982). To generate a similar lure item (*f_j_′* ) for a given target item (*f_j_*), we used the following equation (Murdock, 1995):

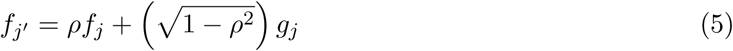

where *ρ* indicates the desired similarity between the target item and the similar lure item and *g_j_* represents an independent random vector. Here, the expected value of the dot product (i.e., similarity) between a target item and the resultant similar lure is *ρ*. To model the learning phase of the item-only version of TODAM, we calculated the memory vector with the following equation (Murdock, 1995; pg. 105 from Kahana, 2012):

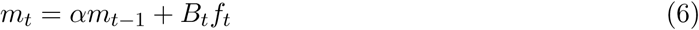

where *α* is a forgetting parameter (i.e., the opposite of a retention parameter, where 0 indicates complete forgetting of previous memories and 1 indicates that the new memory is added to the memory vector from the previous trial without any forgetting), *m_t−_*_1_ is the memory vector from the previous trial, *B_t_* is a vector indicating which features should be learned (more information below), and *f_t_* is the item that is presented at time *t* in the encoding phase of the task. *B_t_* is generated by drawing from the binomial function in numpy with the probability of encoding a feature equal to *p* (i.e., the learning rate of the model) and not encoded with probability 1 - *p*. Therefore, *p* in TODAM is isomorphic to *L* in MINERVA 2. In our current simulations, we set *α* to be equal to 1 because we were not modeling the effects of list position effects and this makes the instantiation more similar to MINERVA 2.

During the test phase, we calculated the global match, *g*, of a probe item to the memory vector using the dot product (using the dot function in numpy):

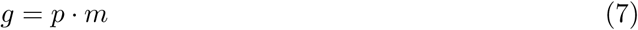

where *p* is the probe item and *m* is the memory vector. MINERVA 2 uses a cube function to generate a nonlinear mapping between the match and the summed similarity; however, TODAM uses a linear similarity function. Also, because TODAM assumes that memory is stored in a single memory vector, the calculation of the similarity is naturally a measure of summed similarity.

For both the MINERVA 2 and TODAM, we simulated forced-choice performance and old/new performance using the general methods described below (see *Simulations of Forced-Choice Perfor- mance*).

#### The Hopfield/Autoassociative Neural Network

We also modeled performance using a Hopfield/autoassociative neural network (Hopfield, 1982; Rizzuto & Kahana, 2001) within Python. Hopfield/autoassociative neural networks are a fully connected, single-layer recurrent neural network that shares functional overlap with the structure and function of the CA3 subregion of the hippocampus (cf. Rolls, 2007; Marr, 1971; McNaughton & Morris, 1987). For example, the majority of connections in CA3 come from the recurrent collaterals (i.e., inputs from other CA3 neurons). For example, Wilson, Gallagher, Eichenbaum, and Tanila (2006) estimated that more than 95% of the inputs to CA3 neurons are recurrent collaterals. Therefore, while this model is still a form of an abstract neural network (i.e., it is not specifically designed to model the anatomical and functional properties of the brain), it bears resemblance to the anatomical, functional, and computational properties of CA3.

Items are also represented as vectors in the Hopfield/autoassociative neural networks. Specifically, we generated items with the features of -1 and 1, with an equal probability of either feature. These values can be thought of as modeling activity levels that are lower, -1, or higher, 1, than the baseline or average activity (pg. 165-166 from Kahana, 2012). To generate similar lure items, we followed a similar procedure to our approach in MINERVA 2. Specifically, we randomly redrew features (i.e., -1 and 1) for a specified proportion of the features using the binomial function from numpy to select the features to be changed.

To simulate learning, we used a probabilistic Hebbian learning paradigm (Rizzuto & Kahana, 2001). The first part of the learning model was the standard autoassociative Hebbian learning rule, which is implemented as the sum of the outer products across all of the trials (cf. Rizzuto & Kahana, 2001):

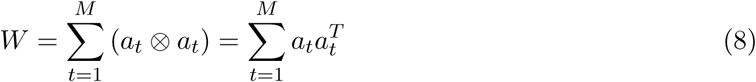

where *W* is the weight matrix, *a_t_* is the activity vector at time t (a column vector), ⊗ represents the outer product (here, using the outer function in numpy), and *a^T^* indicates the transpose of a (i.e., a row vector). Note that following Hopfield (1982) and Kahana (2012, pg. 166) we set the connection between a unit and itself (i.e., the diagonal of the weight matrix) to be equal to 0. We then implemented a probabilistic learning rule by allowing the network to learn each feature (i.e., each connection between simulated neurons/each entry in the weight matrix) with probability *γ* and to not learn each feature with probability 1 − *γ*, which we calculated separately on each trial (Rizzuto & Kahana, 2001). We calculated *γ* using draws from a truncated normal distribution, which allows us to incorporate encoding variability into the encoding phase. Specifically, we generated the probabilities from a normal distribution with a mean of 0.4 and a standard deviation of 0.1 using the function numpy.random.normal. Whenever the draw from the random distribution returned a value that was less than 0 or greater than 1, we redrew from the normal distribution (i.e., this is the truncated aspect of the truncated normal distribution; Rizzuto & Kahana, 2001). Note that Rizzuto and Kahana (2001) modeled different versions of the probabilistic encoding rule; e.g., to allow for symmetric vs. asymmetric learning of associations between items in an associative memory task. However, in our case, we implemented a fully symmetric learning rule. Once we calculated *γ* for a given trial, we then used this value to generate the features to encode via the binomial function in numpy. Specifically, we generated a binomial vector of values with a probability of success equal to *γ*, with a length equal to (*N* ×(*N* − 1))*/*2; i.e., corresponding to the upper triangle of the change-in-weights matrix: Δ. We then placed these features in the upper triangle of the binary change-in-weights matrix (which was initialized as all zeros) and then we made the matrix symmetric with the following equation:

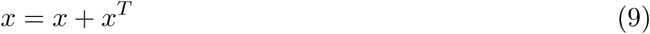

Altogether, the weight changes for the probabilistic Hebbian learning rule for each trial were given by the element-wise multiplication of the change-in-weights matrix with the outer product of the item vectors, which we then summed across all trials, as shown by following equation (cf. Rizzuto & Kahana, 2001):

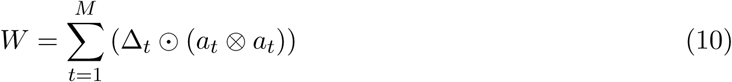

where Δ*_t_* represents a binary change-in-weights matrix (on trial *t*) that implements the probabilistic learning rule described above (values of 1 for weights that will change and 0 otherwise; note that the *t* subscript indicates that the matrix representing the weights to change is trial unique), ⊙ represents the Hadamard product (i.e., the element-wise product) of the two resultant matrices, *a_t_* is the activity vector at time *t*, and ⊗ represents the outer product (here, using the outer function in numpy). As we mentioned above, we set the connection between a unit and itself (i.e., the diagonal of the weight matrix) to be equal to 0 (see Hopfield, 1982 and pg. 166 from Kahana, 2012).

To model the test phase, we used two approaches, one based on a familiarity-based model (for a similar approach to modeling memory within the perirhinal cortex see Bogacz, 1999; Bogacz, Brown, & Giraud-Carrier, 2001) and another based on a recall-based model (for a similar approach to modeling the hippocampus see Norman & O’Reilly, 2003). For the familiarity-based model, we calculated the harmony (*H*, i.e., the match) of the probe item, *p*, to the memory matrix, *W*, using the harmony function (cf. Hopfield, 1982; Smolensky, 1986; see pg. 107 from O’Reilly & Munakata, 2000):

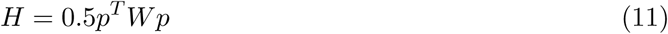

where *p* represents the probe vector (a column vector), W represents the weight matrix, and *p^T^*represents the transpose of the probe vector (i.e., a row vector). Specifically, we used the dot function within numpy for the calculation (0.5 * np.dot(np.dot(p, W), p))). Note that Bogacz (1999) and Bogacz et al. (2001) applied a similar approach to modeling familiarity-based discriminations in the perirhinal cortex (i.e., using the energy of the network, *E*, which is simply the negative value of the harmony of the network: *E* = −*H*).

For the recall-based model, we used Hopfield’s (1982) dynamical rule for updating the trace. Specifically, for each iteration, we calculated the output of the network with the following equation (Hopfield, 1982; Rizzuto & Kahana, 2001; Kahana, 2012, pg. 170):

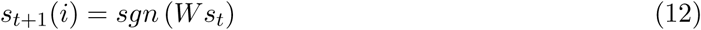

where *sgn* is the sign function (here, set to +1 if the activity of the unit is greater than 0, to -1 if the activity of the unit is less than 0, and randomly to +1 or -1 if the activity of the unit is equal to 0), *W* represents the weight matrix, and *s_t_* represents the trace at time *t*. The initial value of *s* at the start of a test trial (i.e., *s*_0_) was one of the probes (i.e., either a target, a similar lure, or a novel foil item). We used the dynamical retrieval rule, in which we updated the activity of a single unit per iteration (i.e., time step: t, denoted as *s_t_*_+1_(*i*), where the *i*th unit is updated; Hopfield, 1982; Rizzuto & Kahana, 2001). Here, we randomly selected one of the units that did not have the same value as the previous time step. We allowed the network to iterate until it reached a match criterion, *θ*, to one of the items in the training lexicon (here, *θ* = cosine similarity *>* 0.99) or for a maximum number of iterations (*Imax* = 800; Rizzuto & Kahana, 2001). After each trial ended (i.e., either exceeding *θ* or reaching *Imax*), we then calculated the similarity between the probe and the retrieved pattern (cf. Norman & O’Reilly, 2003) using the cosine similarity (i.e., values close to 1 indicate stronger match between the probe item and the recalled item, whereas values closer to 0 indicate a weaker match between the probe item and the recalled item).

For both the familiarity-based model and the recall-based model, we simulated forced-choice performance and old/new performance using the general methods described below (see *Simulations of Forced-Choice Performance*).

#### Simulations of Forced-Choice Performance

To simulate performance on the forced-choice test formats, we calculated the relevant memory scores for each model. Then, we simulated the decision on the forced-choice formats using the following equation (Huffman & Stark, 2017; cf. Hintzman, 1988):

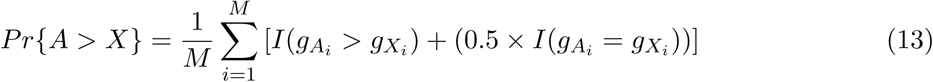

where *M* represents the list length, *I*() is the indicator function that returns the value 1 if the statement is true and 0 otherwise, *g_Ai_* is the memory score for the target item on trial i, and *g_xi_* is the memory score for the distractor item (i.e., a novel foil or a similar lure) on trial i. Note that the second part of the equation deals with guessing by simulating that the model would get 0.5 of the guess trials correct (i.e., the model would randomly guess when the memory score for the target and distractor items match). We could have also simulated “tie-breaking” by randomly selecting an item on a trial-by-trial basis (cf. Norman & O’Reilly, 2003); however, the equation above will produce the same results on average.

#### Simulations of Old/New Performance

To simulate performance on the old/new test formats, we calculated the relevant memory scores for each model. Then, we simulated the old/new test formats with confidence ratings by calculating the proportion of hits and false alarms at several values of the memory strength—i.e., we modeled the confidence ratings as increasing values of the memory scores. Specifically, here we used 10 confidence bins by linearly spacing from the minimum value of the target memory score (i.e., the lowest value that would induce any hits) to the maximum value of the distractor memory score (i.e., the highest value that would induce any false alarms). We then calculated the simulated hit rates and false alarm rates by calculating the number of responses within each simulated confidence bin: i.e., the number of values that are greater than or equal to the minimum cutoff and less than the cutoff of the next confidence bin. Note that for the weakest memory condition, we also added all of the distractor responses that were less than the minimum cutoff (i.e., the lowest memory score for the target distribution) and for the strongest memory condition, we added all of the target responses that were greater than the maximum cutoff (i.e., the highest memory score for the distractor distribution). Note, we calculated the minimum/maximum confidence bins in this manner (e.g., rather than allowing a nonzero hit rate with a zero false alarm rate) because the binormal model that we used for calculating performance assumes that the hit rate/false alarm rate plot (i.e., the receiving operator characteristic curve) passes through the origin: i.e., (0, 0). We then calculated the cumulative proportions of hits and false alarms based on these simulated confidence bins (from the “most confident old”/“least confident new” to the “least confident old”/“most confident new”). Then, we calculated our measure of performance, the area under the receiver operating characteristic (ROC) curve (*A_z_*), using the same techniques as we used for our empirical data (see *Comparison of the Forced-Choice and Old/New Test Formats* for more details and rationale of using this measure).

### Results and Discussion

#### Predictions of Forced-Choice Performance

As we discussed in the Introduction, the CLS model predicts that the hippocampus will exhibit similar performance on the A-A’ and the A-B’ forced-choice test formats (see Figure 2a and Norman & O’Reilly, 2003). In fact, Norman and O’Reilly (2003) reported that there are even times in which the hippocampus performs better on the A-B’ than the A-A’ test format because the hippocampus employs recall-to-reject in their model, thus the A-B’ test format gives the hippocampus two chances to recall-to-reject the stimulus, thus any benefit of covariance in the A-A’ is offset (or even overcompensated for) by the additional opportunity to recall-to-reject a given stimulus. In contrast, the global matching models, MINERVA 2 (see Figure 5a) and TODAM (see Figure 6a), predict a strong A-A’ *>* A-B’ test format effect (Hintzman, 1988; Huffman & Stark, 2017). Moreover, we found that both the “familiarity” (see Figure 7a) and the “recall” version (see Figure 8a) of the Hopfield/autoassociative neural network predicted a similar A-A’ *>* A-B’ test format effect. Therefore, in summary, MINERVA 2, TODAM, and both versions of the Hopfield/autoassociative neural network suggest that there will be a strong A-X *>* A-A’ *>* A-B’ test format effect (also see the neocortical model from Simulation 4 in Norman & O’Reilly, 2003), which differs from the CLS hippocampal model, which predicts a null A-A’ *>* A-B’ test format effect (see Norman & O’Reilly, 2003; Migo et al., 2009). As we report below, we found a clear effect of test format, thus suggesting that human memory performance is primarily driven by the distributed representations of the neocortex rather than the pattern-separated representations of the CLS hippocampal model.

#### Predictions of the Transfer Function between Performance and Target-Lure Similarity

As we discussed in the Introduction, the CLS model predicts that the hippocampus would not exhibit a clear and continuous transfer function between simulated performance and targetlure similarity. Specifically, the transfer function between performance and target-lure similarity is flat or null until only very high levels of similarity (i.e., *>* 0.875; see Figure 2b and Norman & O’Reilly, 2003). In contrast, the CLS model of the neocortex exhibits a much more continuous effect of target-lure similarity on simulated performance (Norman & O’Reilly, 2003). In general, all of the distributed memory models predict a relatively continuous transfer function between memory performance and target-lure similarity (MINERVA 2: Figure 5b; TODAM: Fig- ure 6b, the familiarity-based Hopfield/autoassociative neural network: Figure 7b; the recall-based Hopfield/autoassociative neural network: Figure 8b). Therefore, we aimed to test these competing predictions to determine whether human behavioral performance is primarily driven by the pattern-separated representations of the hippocampus vs. distributed representations (e.g., the neocortex) vs. a composite representation between these systems. As we reported above (see Quantification of Target/Lure Similarity with an Artificial Neural Network), we found a continuous (and possibly linear) mapping between performance and simulated target-lure similarity, thus supporting the distributed memory models. Moreover, as we discuss below, we replicated this effect in a new sample of participants on the old/new task (see *Old/New Data*) and extended this finding to the forced-choice test formats (see *Forced-Choice Data*).

#### Predictions of the Relationship between Forced-Choice and Old/New Performance

We next aimed to generate predictions about the relationship between performance on various test formats. Norman and O’Reilly (2003) suggested that there are many cases in which hippocampal and neocortical performance will not be correlated, which means that performance across test formats will not always be correlated (e.g., A-A’ performance might not correlate with A-B’ performance). Moreover, as we mentioned above, the CLS hippocampal model predicts that performance will generally be the same across the A-A’, A-B’, and the old/new with targets and similar lures (Norman & O’Reilly, 2003). In contrast, the distributed memory models make clear predictions about the relationship between performance on the various test formats. For example, we modeled individual differences in performance by changing the learning rate in the models. We found that the models predicted clear differences in performance as a function of learning rate (see Figure 9; note: this figure depicts MINERVA 2, but results were similar in the other distributed memory models). Moreover, when we investigated the relationship between performance on the forced-choice vs. old/new test formats, we found that the distributed memory models predicted that performance on the A-X was roughly equivalent to the old/new test format with targets and novel foils (i.e., the regression between performance on the two formats produced a slope of approximately 1; see Figure 10a), performance on the A-A’ test format is predicted to be better than performance on the old/new test format with targets and similar lures (i.e., the regression between performance on the two formats produced a slope that is greater than 1; see Figure 10b), and performance on the A-B’ test format was roughly equivalent to performance on the old/new test format with targets and similar lures (i.e., the regression between performance on the two formats produced a slope of approximately 1; see Figure 10c; also see below, including Figure 15). Additionally, we generated predictions of the similarity of performance on all 5 test formats using a random-effects-style simulation and we found that the distributed memory models suggest that performance will be significantly correlated between all of the test formats (see Figure 11). Thus, while there are conditions in which the CLS model predicts correlated performance between tasks (Norman & O’Reilly, 2003), if we observe 1) roughly equivalent performance between the A-X and old/new test format with targets and novel foils, 2) better performance on the A-A’ test format than both the A-B’ test format and old/new test format with targets and similar lures, 3) roughly equivalent performance between the A-B’ and the old/new test format with targets and similar lures, 4) strong correlations between test formats, it will provide stronger evidence that human memory performance is primarily driven by the distributed representations of the neocortex than the pattern-separated representations of the hippocampus.

**Figure 9:**
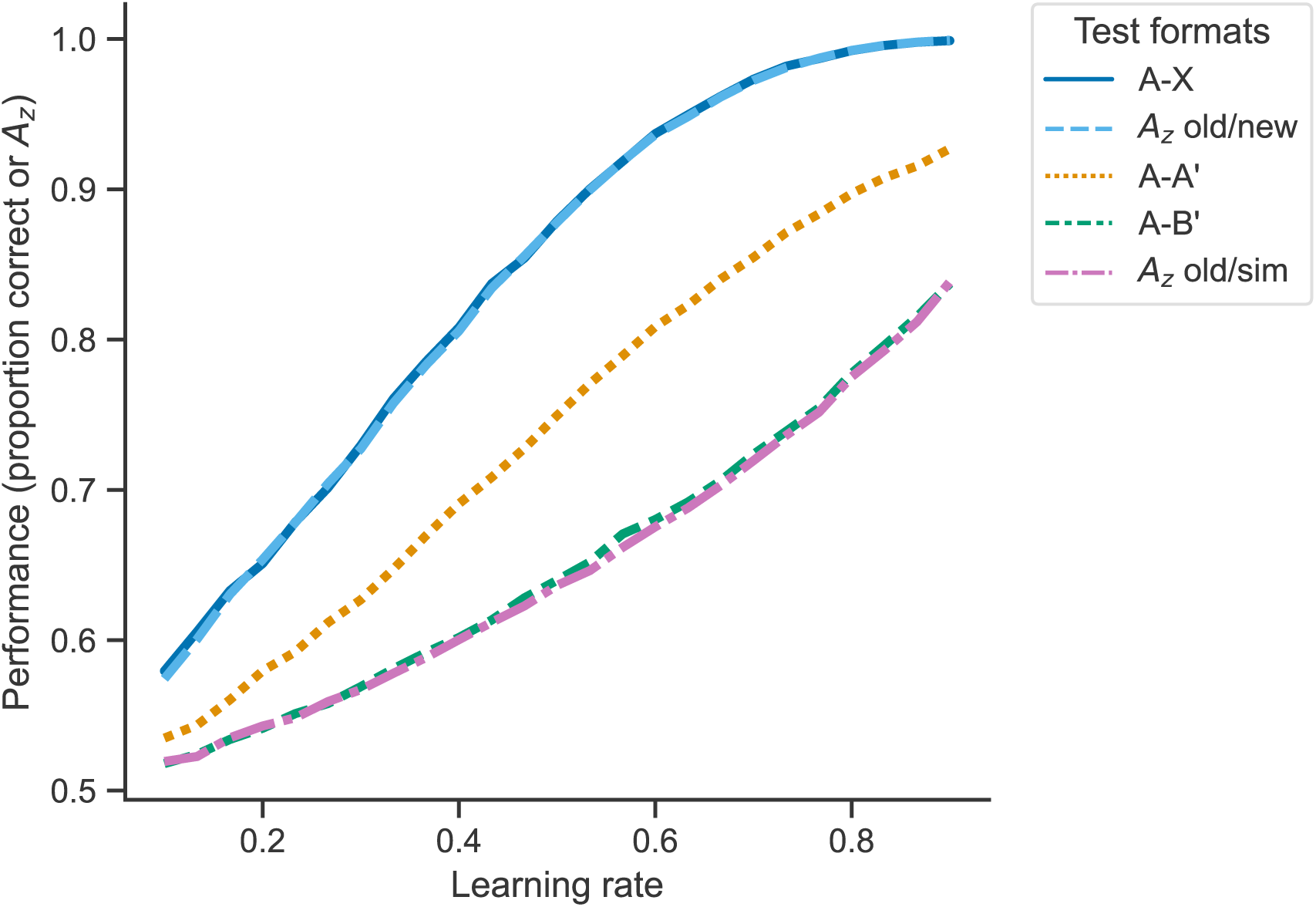
Simulations of individual differences in learning rates reveal performance tuning curves across the 5 test formats. The distributed memory models predict that performance on the A-X and *A_z_* for targets vs. novel foils will be roughly equivalent as will performance on the A-B’ and the *A_z_* for targets vs. similar lures, while performance on the A-A’ test format will be intermediate. Here, we show the results of MINERVA 2, but the other distributed memory models made very similar predictions.

**Figure 10:**
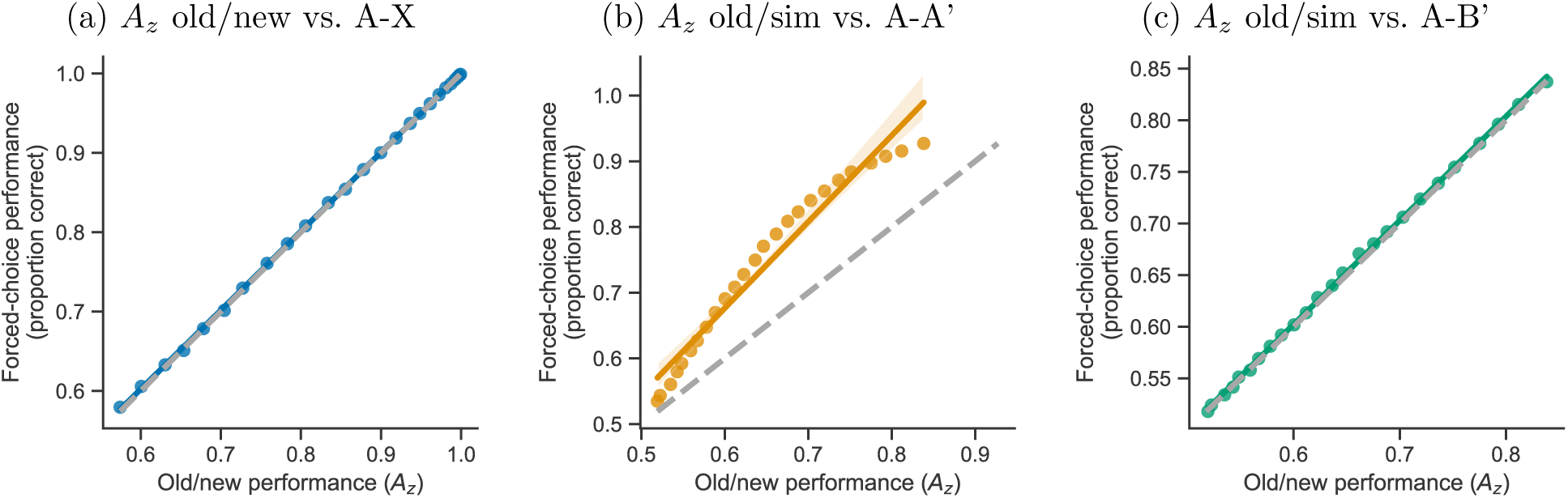
The distributed memory models make clear predictions about the relationship of performance on the forced-choice and old/new test formats. (a) The models predict that performance on the A-X and the old/new test format with targets and novel foils will be approximately equivalent (i.e., a slope of approximately 1). (b) The models predict that performance on the A-A’ test format will be significantly better but also strongly related to performance on the old/new test format with targets and similar lures (i.e., a slope that is greater than 1). (c) The models predict that performance on the A-B’ test format will be approximately equivalent to performance on the old/new test format with targets and similar lures (i.e., a slope of approximately 1). Please compare these results to Figure 14, Figure 15, and Figure 16.

**Figure 11:**
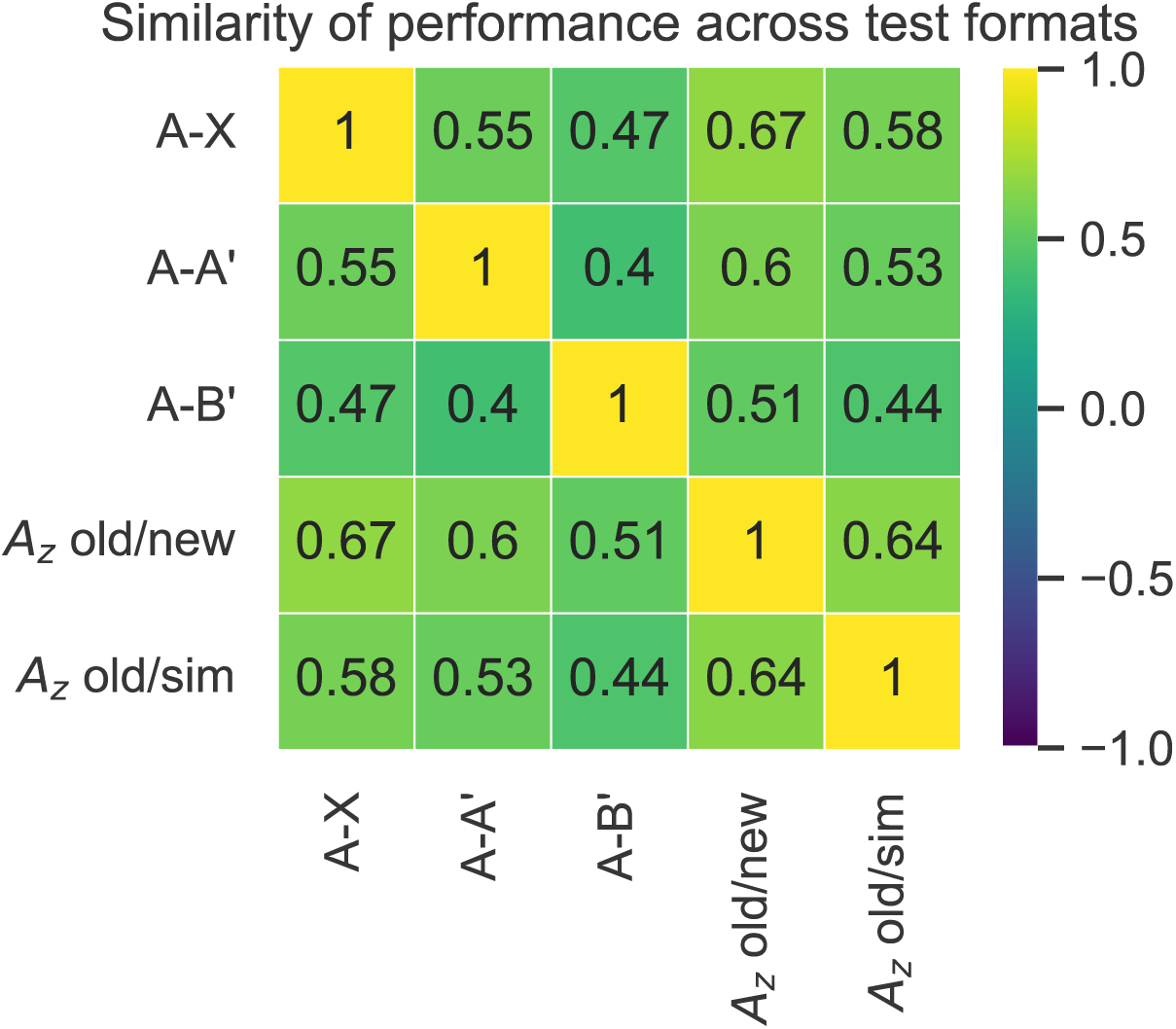
The distributed memory models predict that individual differences in learning rates will lead to correlated performance between all 5 test formats. The correlation matrix depicts simulated performance in MINERVA 2, but all of the models made similar predictions. Please compare these results to the empirical results in Figure 17.

**Figure 12:**
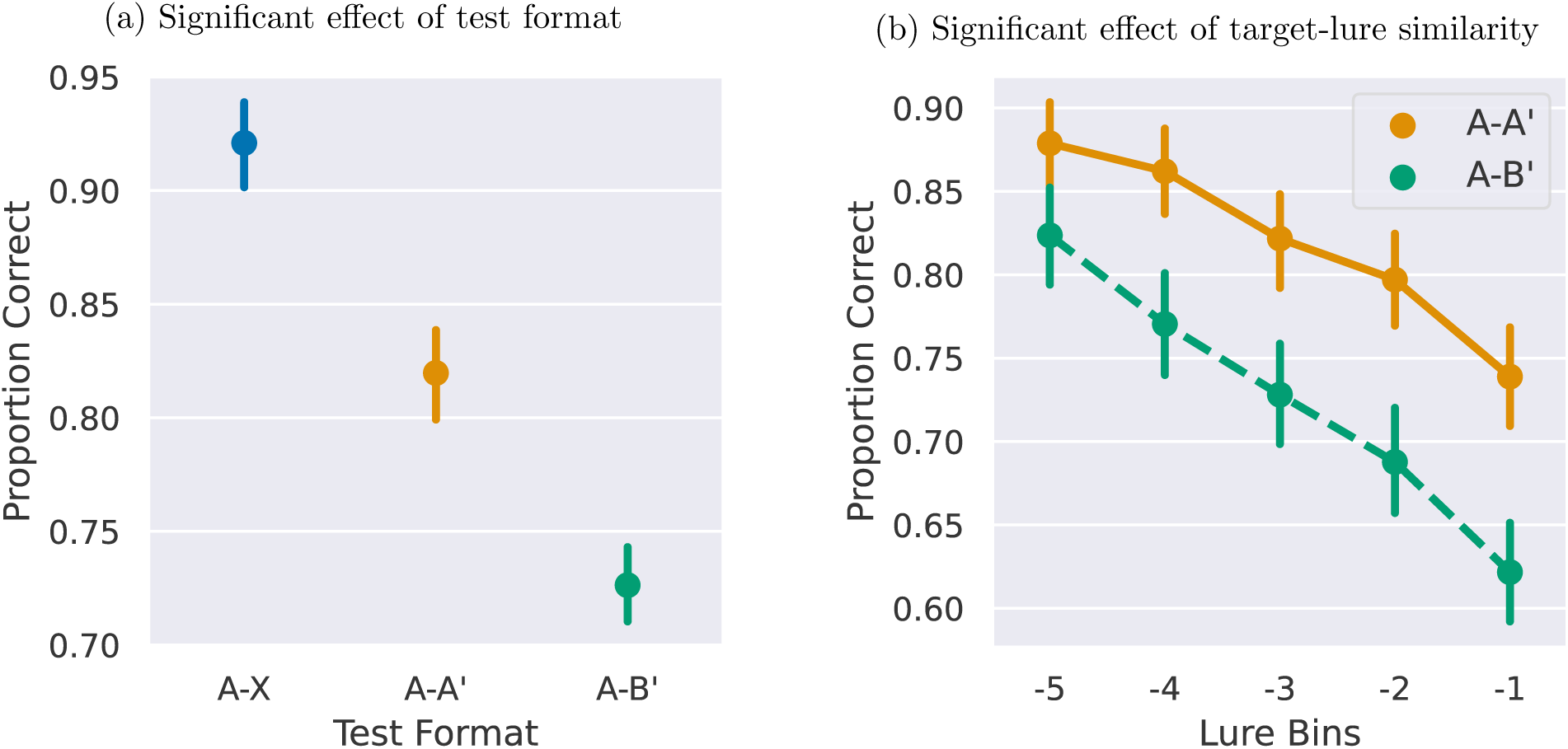
Human performance on the forced-choice test formats differed as a function of test format and as a function of target-lure similarity. (a) We observed a significant effect of test format: participants performed best on the A-X format followed by the A-A’ format followed by the A-B’ format, which is consistent with the distributed memory models (but not the CLS hippocampal model). (b) We observed a significant effect of lure bin (i.e., mnemonic similarity from previous old/similar/new experiments) on performance of the A-A’ and A-B’ test formats. Moreover, the lure-bin effect was remarkably similar between conditions, although participants performed significantly better on the A-A’ than the A-B’ format. Note: for visualization, the data are plotted here as the proportion correct across all trials; however, we analyzed our data using generalized linear models with a binomial family. Additionally, for visualization, we multiplied the lure bins by -1 so that the least similar bins are to the left of the plot and the most similar bins are to the right of the plot, thus enabling an easier comparison with the predictions of the computational models. Altogether, these results support the predictions of the distributed memory models (see Figure 5, Figure 6, Figure 7, and Figure 8) but are at odds with the predictions of the CLS hippocampal model (see Figure 2). Points represent the mean and the error bars indicate the 95% confidence intervals.

**Figure 13:**
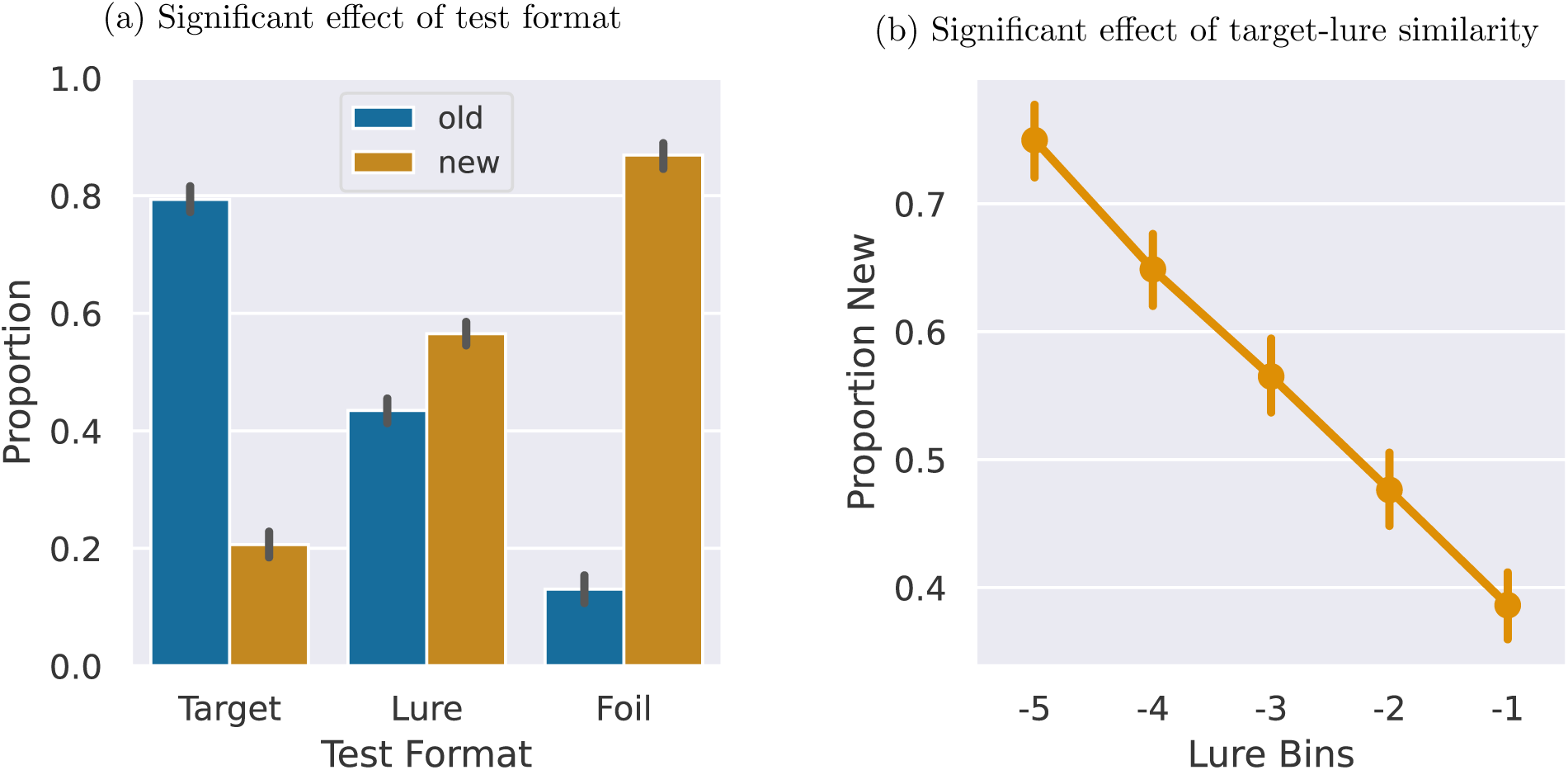
Human performance on the old/new format differed as a function of test format and as a function of target-lure similarity. (a) We observed a significant effect of test format: participants made a different number of “old” and “new” responses across the targets, similar lures, and novel foils. (b) We observed a significant effect of lure bin (i.e., mnemonic similarity from previous old/similar/new experiments) on the number of times participants responded “old” to similar lures. Note: for visualization, the data are plotted here as the proportion correct across all trials; however, we analyzed our data using generalized linear models with a binomial family. Additionally, for visualization, we multiplied the lure bins by -1 so that the least similar bins are to the left of the plot and the most similar bins are to the right of the plot, thus enabling an easier comparison with the predictions of the computational models. Altogether, these results support the predictions of the distributed memory models (see Figure 5, Figure 6, Figure 7, and Figure 8) but are at odds with the predictions of the CLS hippocampal model (see Figure 2). Bars and points represent the mean and the error bars indicate the 95% confidence intervals.

**Figure 14:**
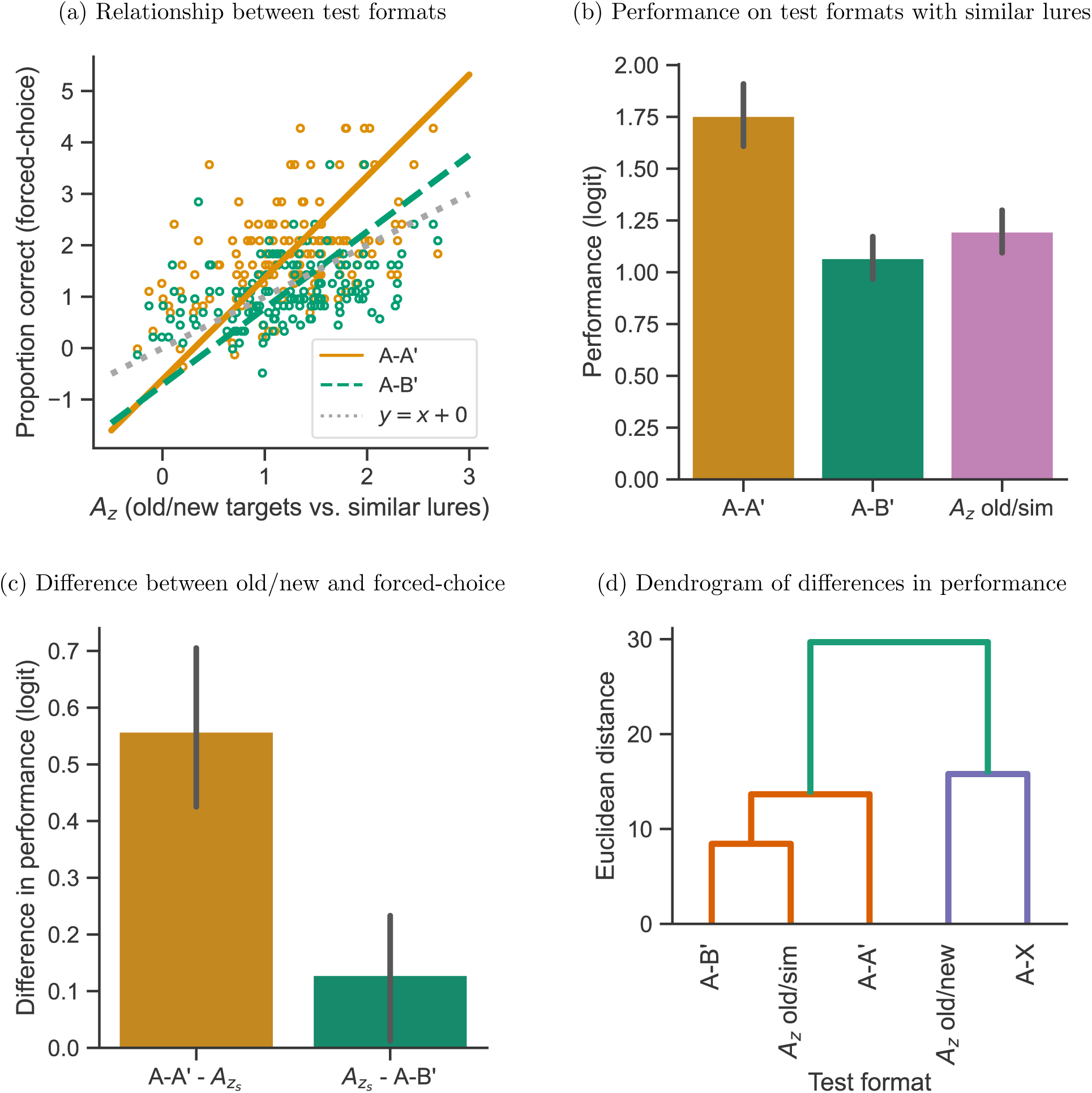
Evidence that human performance on the old/new test format with targets and similar lures is most similar to the A-B’ test format. (a) The comparison between the old/new test format with targets and similar lures vs. the A-A’ and A-B’ test formats. Note that the comparison with the A-B’ test format produced the closest slope to 1. (b) The logit-transformed performance revealed that (c) performance was most similar between the A-B’ test format and the old/new test format with targets and similar lures. (d) Dendrogram analysis also revealed clustering of the test formats. Altogether, these results support the predictions of the distributed memory models (see Figure 9, Figure 10, and Figure 15). Error bars represent the 95% confidence intervals.

**Figure 15:**
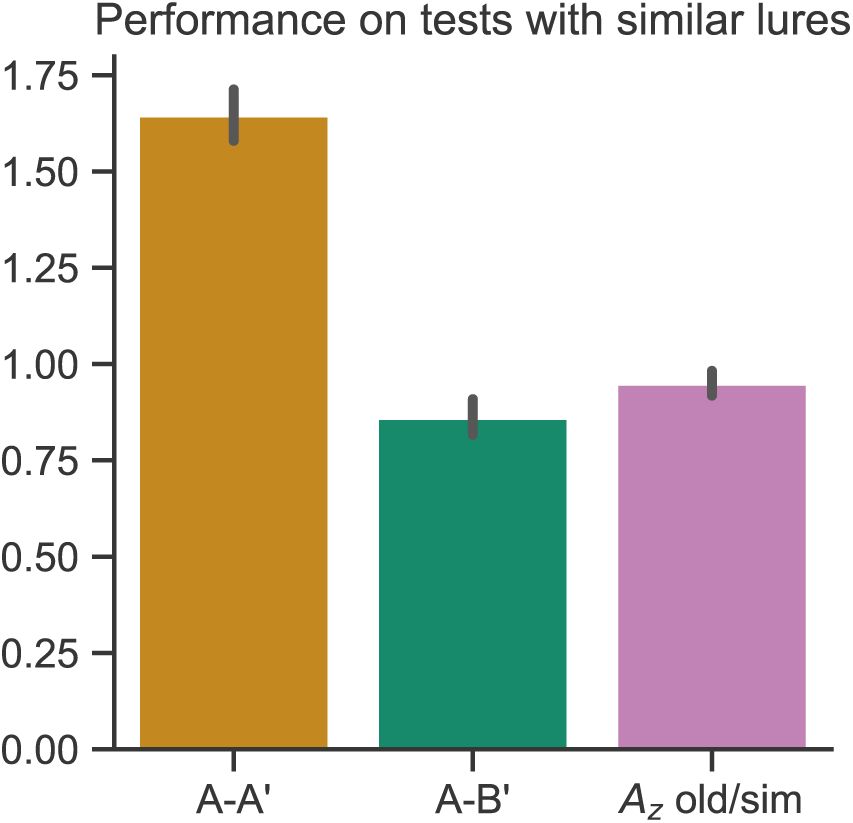
A follow-up, random-effects-style analysis of the distributed memory models revealed the same findings as the empirical data (see Figure 14b). This plot is from MINERVA 2, but the other distributed memory modes made the same predictions. Bars represent the mean and the error bars indicate the 95% confidence intervals.

## Behavioral Experiment

Our final overarching goal was to collect a large sample of data from human participants to test the predictions from the computational models. We were specifically interested in determining whether human memory performance is primarily driven by the pattern-separated representations of the hippocampus (e.g., dentate gyrus) vs. distributed representations (e.g., the neocortex or hippocampal subregions CA1 or CA3) vs. a composite representation across these systems.

### Methods

#### Participants

Participants consisted of 153 undergraduate students at Colby College. The mean age of participants was 19.5 years old, with a standard deviation of 1.17 and a range between 18 and 22. 108 participants identified as female, 42 participants identified as male, 2 participants identified as non-binary, and 1 participant used another term or option not listed. Their participation lasted no longer than 60 minutes, and all participants consented to participation in accordance with the Institutional Review Board at Colby College. Participants were recruited within the Colby Psychology Department, and they received course credit for their participation. We planned to collect data from at least 100 participants given that some of our measures included studying individual differences and correlations between performance on different test formats. To ensure the success of the Receiver operating characteristic (ROC) analysis (see below), we excluded 5 participants whose answers were not distributed across confidence bins (e.g., if they only used two confidence bins). We also excluded 3 participants whose responses in the encoding phase were biased (i.e., if they selected 10 times more ‘outdoor’ than ‘indoor’ or vice versa, or if they did not respond to more than 100 trials) and the correct rate in all test formats of the forced-choice version were less than 0.7. The final analysis included 145 participants.

#### Behavioral Task Design

The Mnemonic Similarity Task (MST; Kirwan & Stark, 2007; Stark et al., 2013, 2015, 2019; Huffman & Stark, 2017) has been developed to study memory performance for targets, similar lures, and novel foils. The MST has been employed with several versions of the test format. The typical versions of the task include an Old/Similar/New test format, in which participants are instructed to respond “old” if they think that they have previously seen the exact item, “similar” if they think that they have previously seen a similar item, and “new” if they think that they did not see this item or an item like it in the encoding phase. Other studies have used an Old/New test format, in which participants are instructed to respond “old” if they believe that they saw the exact item and to respond “new” otherwise (i.e., to similar lures and novel foils; cf. the “veridical” version of the task in Experiment 4 in Stark et al., 2015). Moreover, studies using the Old/New test format have also occasionally included confidence ratings, which allows for analysis with methods from signal detection theory, including the receiver operating characteristic (ROC) curve (cf. Stark et al., 2015; Loiotile & Courtney, 2015). Finally, previous studies have also studied performance on a forced-choice version of the task (e.g., Huffman & Stark, 2017; Rollins et al., 2019; cf. Jeneson et al., 2010; Tulving, 1981). Specifically, participants in these experiments were tested under three forced-choice formats: A-X (a target and an unrelated, novel foil; traditional forced-choice recognition), A-A’ (a target and a corresponding similar lure; e.g., a picture of an apple and a similar apple), and A-B’ (a target and a noncorresponding similar lure; e.g., a picture of an apple and a basketball that is similar to one that was studied in the encoding phase; note, we use the nomenclature of Tulving, 1981). Therefore, these various test formats allow us to test how the test format itself influences performance on the task, which we explore here.

Our experiment consisted of two versions of the Mnemonic Similarity Task (forced-choice and old/new with confidence ratings), one automatic 10-minute rest between the two versions, and questionnaires at the end. We ran our experiments online within the cognition.run platform. After the welcome and introduction messages, the experiment website randomly assigned participants to either begin with the forced-choice version of MST or the old/new version. In both versions, participants first performed an encoding task in which they indicated whether each presented image depicted more of an indoor or an outdoor item (i.e., the indoor/outdoor task, which is an incidental encoding task; see Figure 1). Each encoding phase consisted of 140 images, which were displayed for 2,000 ms with a 500-ms interstimulus interval.

In the test phase of the forced-choice MST (see Figure 1), participants performed a memory test that contained three test formats: 1) A-X (i.e., a target and a novel foil), 2) A-A’ (i.e., a target and a corresponding similar lure), and 3) A-B’ (i.e., a target and a noncorresponding lure object from a different pair), as in our previous research (Huffman & Stark, 2017). Each test format included 35 trials. We matched the “lure bin” (i.e., similarity level) of the target and lure/foil item in each pair of stimuli to account for any potential effects of encoding difficulty of the stimuli (as in Huffman & Stark, 2017). The test formats were presented in a random order on a trial-by-trial basis. Every trial contained two objects, one on the left side of the screen and one on the right side, and the target was randomly assigned to one side on each trial. Participants were instructed to select the exact object they saw during the encoding phase via button press. The images were displayed either until the participant made a response or for 4 s, after which the image disappeared but the participants had an unlimited response window.

In the test phase of the old/new MST, participants performed a memory test that contained three types of objects: 1) targets (i.e., exact repetitions of images they saw during the indoor/outdoor task), 2) similar lures (i.e., images that were similar to one they saw during the indoor/outdoor task), 3) novel foils (i.e., new images that were not in a similar image pair to one they saw during the indoor/outdoor task). The task included 70 targets, 70 similar lures, and 52 novel foil trials (i.e., the remainder of the number of available stimuli because each MST image set has 192 images). On each test trial, one object was presented in the center of the screen, and participants were instructed to identify whether they saw it during the encoding phase (i.e., “old” object) or not (i.e., “new” object) via button press. We instructed participants to respond “old” if they saw this exact image during the indoor/outdoor task (i.e., they should respond “old” only to targets) and to respond “new” otherwise (i.e., to novel foils and to similar lures). The images were displayed until participants made a response or for 2 s, after which the image disappeared and left an unlimited response window. Following the old/new response, participants indicated their confidence in their decision, with the following options: “very sure”, “somewhat sure”, “somewhat unsure”, “very unsure”. As we describe below, we were able to convert the two responses (i.e., old, new) and the four confidence bins to a total of 8 possible responses (i.e., 2 × 4) for the ROC analysis.

After the MST, participants completed questionnaires about their demographic information, video game and GPS use, autobiographical memory, sleepiness, and COVID-19 information. Specifically, we used the Survey of Autobiographical Memory (SAM; Palombo, Williams, Abdi, & Levine, 2013) to assess participants’ trait mnemonics and Stanford Sleepiness Scale (SSS; Hoddes, Zarcone, Smythe, Phillips, & Dement, 1973) to assess participants’ sleeping performance. We did not analyze the questionnaires here.

#### Analytic Methods for the Forced-Choice Data

To analyze the forced-choice data, we used generalized linear models with the lme4 package in R (version: 1.1-21; Bates, Mäachler, Bolker, & Walker, 2015). Specifically, we used the glmer function with a binomial family because our data were binomial (i.e., on each trial a participant either selected the correct answer or the incorrect answer). We fit our models with random intercepts for the participants. We assessed significance of the effects using a likelihood ratio test, using the anova function in R with the test set to “LRT”. We also report the Bayesian information criterion (BIC), where lower values indicate better model fits while penalizing for model complexity. To test whether there was a significant effect of test format, as in previous papers (Huffman & Stark, 2017), we compared a model with the effect of test format to a random intercepts and slopes model:

Random-intercepts-and-slopes model:

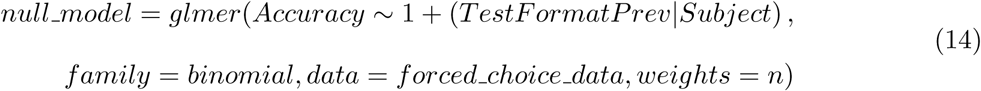

Test-format model:

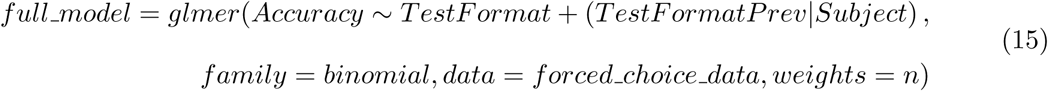

To test whether the empirical data fit better with the predictions of the distributed memory models (i.e., A-X *>* A-A’ *>* A-B’) than the CLS hippocampal model (i.e., A-X *>* A-A’ = A-B’; a null effect of the A-A’ and A-B’ test formats), we compared the fit of a model with the test formats equal to the mean performance from Experiment 2 in Huffman and Stark (2017) (A-X: logit(0.959) A-A’: logit(0.836), A-B’: logit(0.756)) vs. a model with equal performance on the A-A’ and A-B’ test (A-A’ = A-B’: mean(logit((0.836, 0.756)))). Specifically, we ran Equation (15) two times, once with the previous performance values and once with the CLS hippocampal model predictions (i.e., the *TestFormat* values differed for each run of the model). To keep the models consistent in terms of the random effects, in both models the random effects structure included the random effects data from the previous performance values (i.e., termed *TestFormatPrev* above). We then compared the Bayesian information criteria (BIC) between these two models (i.e., because zero parameters differed), by comparing both the raw BIC values (i.e., lower BIC indicated the preferred model) as well as the approximation of Bayes factors using Equation (2) (Wagenmakers, 2007; Jarosz & Wiley, 2014). To test whether there was a significant effect of lure bins, as in previous papers (e.g., for old/similar/new test format: Stark et al., 2013), we included a parameter for the lure bins in our model. Specifically, we compared the test-format model with a test-format-and-lure-bin model.

Random intercepts and slopes (for shortening equations below):

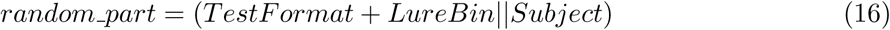

We used a random effects model with random slopes and intercepts for the test format and the lure bin within each subject (and uncorrelated between the slopes and intercepts, hence the || in the equation above). We did not include the random effects for the interaction because the model produced a singular fit and failed to converge. Thus, we chose the maximal model that converged (Barr, Levy, Scheepers, & Tily, 2013; Bates, Kliegl, Vasishth, & Baayen, 2015); moreover, as we discuss in the Results, the interaction model was not significant anyway, thus the random effects model in no way limits our conclusions here. We then compared several models to determine how well each model fit the data.

Random-intercepts-and-slopes model:

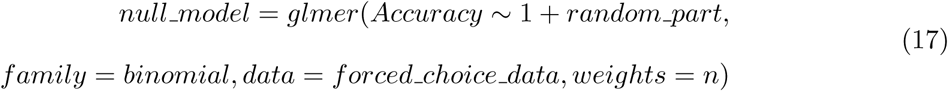

Test-format model (for comparison to Equation (19) with the A-A’ and A-B’ data):

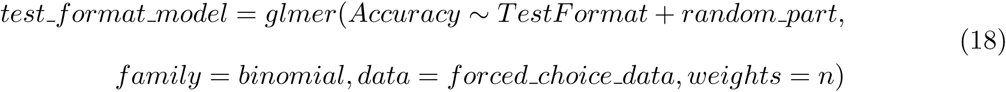

Test-format-and-lure-bin model:

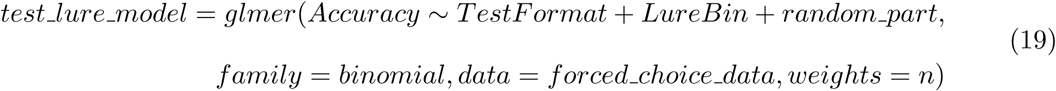

Here, we focused our analysis on the A-A’ and A-B’ test formats since the A-X test format did not manipulate the similarity to previously viewed items. We also investigated whether the effect of lure bins differed as a function of test format by comparing a test-format by lure-bin interaction model with the test-format-and-lure-bin model.

Test-format by lure-bin interaction model:

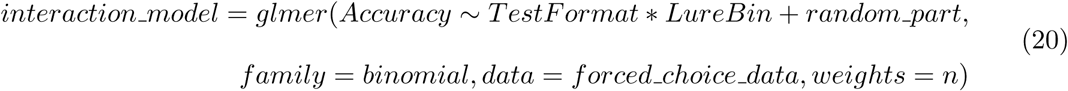

Note, for these analyses, we normalized our lure bins (range: [1..5]) so that they center around 0 by subtracting 3 from each lure bin (range: [−2..2]). The lure bins are organized from the most similar (lowest lure bin) to the least similar (highest lure bin; see Figure 1), thus a positive beta estimate from these models would be indicative of a positive monotonic relationship between empirical mnemonic similarity and performance on the task.

We also followed up these analyses by comparing the following lure-bin model with an intercept- only version of the model (i.e., replacing *LureBin* with 1 in Equation (21)), separately for each condition (i.e., the A-A’ and A-B’ test format data).

Lure-bin model (for testing within a single test format):

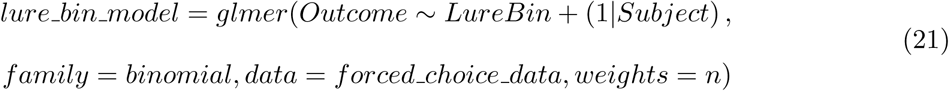

#### Analytic Methods for the Old/New Data

To test whether there was a significant effect of lure bins on performance of the old/new test format with targets and similar lures, as in previous papers (e.g., for old/similar/new test format; Stark et al., 2019), we included a parameter for the lure bins in our model. First, we converted the participants’ responses from “old” and “new” to binary responses, 0 if they selected “old” (i.e., the incorrect response) and 1 if they selected “new” (i.e., the correct response). Then we compared a lure-bin model (zero-meaned as in our forced-choice analysis; see above) to an intercept-only model using a binomial model as we did for our forced-choice data. Note that the more maximal model did not produce a singular fit here, thus our models were as follows:

Random-intercepts-and-slopes model for the old/new data:

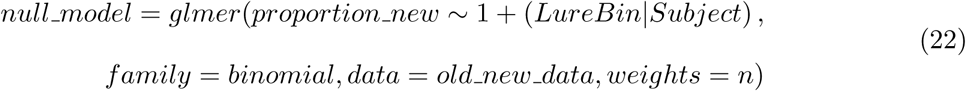

Lure-bin model for the old/new data:

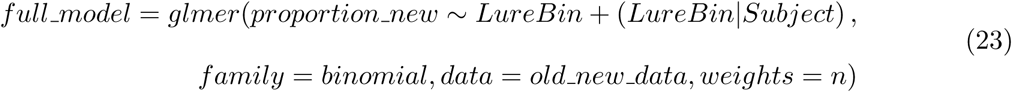

#### Comparison of the Forced-Choice and Old/New Test Formats

One of the key motivations of the current set of experiments was to test the competing predictions from the CLS hippocampal model (Norman & O’Reilly, 2003) vs. the distributed memory models (MINERVA 2, TODAM, and two versions of the Hopfield/autoassociative neural network) about the relationships between performance on the old/new and forced-choice test formats. To allow us to compare the results of the old/new test format to the forced-choice test format, we calculated the area under the receiver operating characteristic (ROC) curve based on the old/new with confidence ratings responses for our various conditions. Previous theories and empirical research have suggested that the area under the ROC curve is mathematically equivalent to proportion correct on a two-alternative forced-choice test format (Green & Moses, 1966; Swets & Pickett, 1982; Stanislaw & Todorov, 1999). Swets and Pickett (1982) and Stanislaw and Todorov (1999) argued that the best measure for calculating the area under the ROC curve is to use maximum likelihood estimation to fit the z-transformed ROC curve with a binormal model, a measure referred to as *A_z_*. Given that the procedure to calculate *A_z_*uses a binormal model, it does not assume that the target and distractor distributions have equal variance (i.e., they can have unequal variance).

We developed our own code for the binormal model within R. The binormal model involves estimating the intercept (*a*), slope (*b*), and estimates of latent cut points within the decision space based on the confidence ratings (the *Z_k_*’s, where the number of cut points is equal to the number of confidence bins - 1). We calculated the initial *a* (intercept) and *b* (slope) using the “unbiased” slope method described by Stanislaw and Todorov (1999). Specifically, we first z-transformed the hit rate and false-alarm rates using the qnorm function in R (a function often referred to as Φ*^−^*^1^ [the “inverse phi” function] in signal detection theory, which converts probabilities to z scores; Stanislaw & Todorov, 1999). Then, we fit two linear regression models: 1) a linear regression with the z-transformed hit rate as the independent variable and the z-transformed false-alarm rate as the dependent variable, 2) we fit another model with the z-transformed false-alarm rate as the independent variable and the z-transformed hit rate as the dependent variable. Then, we calculated the “unbiased” slope by combining the slopes from these two models (Stanislaw & Todorov, 1999):

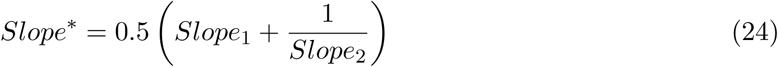

We calculated the initial intercept using the following equation (Stanislaw & Todorov, 1999):

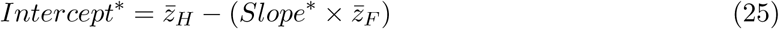

where *z̄_H_* is the mean of the z-transformed hit rates and *z̄_F_* is the mean of the z-transformed false-alarm rates. We then calculated the initial break points (i.e., *Z_k_*) by passing the true negative rates, the initial *x_k_* values, (for the distractor responses; i.e., 1 - false-alarm rates) to the *qnorm* function in R a function often referred to as Φ*^−^*^1^ [the “inverse phi” function] in signal detection theory, which converts probabilities to z scores; Stanislaw & Todorov, 1999). We then used maximum likelihood estimation with a binormal model to optimize the estimates of a, b, and the *Z_k_*values using the following steps. For the distractor distribution (i.e., the novel foils or the similar lures), we calculated the probability that a participant made response *j*, *R_j_*, given stimulus 1, *S*_1_, using the following equation (Dorfman & Alf, 1969):

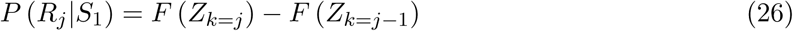

Where *Z_k_* = (*x_k_* − *µ*_1_)*/σ*_1_ (note: these values are then updated in the optimization function described below; see above for explanation of how we calculated the initial *Z_k_*values), and *F* is the cumulative normal distribution (a function often referred to as the Φ [“phi”] function in signal detection theory, which converts z scores into probabilities; Stanislaw & Todorov, 1999), which we calculated with the *pnorm* function in R.

For the target distribution, we calculated the probability that a participant made response *j*, *R_j_*, given stimulus 2, *S*_2_, using the following equation (Dorfman & Alf, 1969):

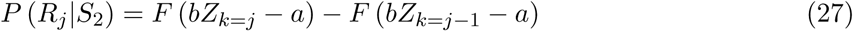

where *b* = *σ*_1_*/σ*_2_, and *a* = (*µ*_2_ − *µ*_1_) */σ*_2_. Then, we maximized the following log-likelihood function (Dorfman & Alf, 1969):

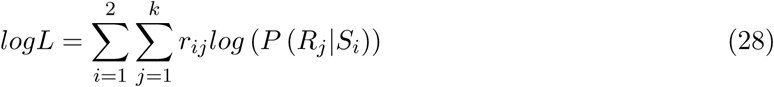

where *r_ij_*represents the number of responses, *R_j_*’s, to stimulus *i* (i.e., either the distractor [*i* = 1] or the target [*i* = 2]), and we used the *optim* function in R with the method set to BFGS (i.e., the Broyden-Fletcher-Goldfarb-Shanno algorithm; Broyden, 1970; Fletcher, 1970; Goldfarb, 1970; Shanno, 1970). Note, the optim function minimizes a given input function, thus we passed the negative value of the log-likelihood function above to the optim function, which, in practice, maximizes the log-likelihood function. Maximizing the log-likelihood function simultaneously optimizes the intercept (*a*), slope (*b*), and the *Z_k_*values. Note, if participants made zero responses at a given confidence level for both the distractor trials and the target trials, then we dropped that bin from the analysis above (i.e., if a participant only used 7 of the possible 8 bins, then k would equal 6 rather than the typical value of 7 [i.e., n - 1 = 8 - 1 = 7]). Additionally, if any bins resulted in a cumulative hit or false alarm rate of 0, we converted it to 0.5 / trials, or if any bins resulted in a cumulative hit or false alarm rate of 1, we converted it to (trials - 0.5) / trials (Dorfman, 1982; Stanislaw & Todorov, 1999).

We then calculated the area under the ROC curve (Az) using the following equation:

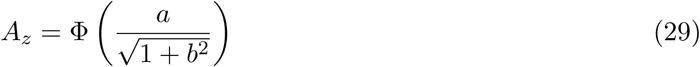

where *a* and *b* are calculated via the binormal model above and Φ is the standard normal cumulative distribution function (calculated with the *pnorm* function in R).

To compare the old/new data to the A-A’ and A-B’ test formats, we calculated *A_z_* for the comparison between targets and similar lures within each participant. Similarly, to compare the old/new data to the A-X test format, we calculated *A_z_* for the comparison between targets and novel foils within each participant.

We then compared the predictions of the distributed memory models by comparing performance between the various test formats. While the *A_z_* measure is directly interpretable as the predicted proportion correct on a two-alternative forced-choice test format, we cannot use binomial models to analyze the data because these are not proportional data (i.e., we would not get integer counts of performance). Therefore, we instead applied the logit function to transform our data for analysis with other methods, as we describe below.

First, we aimed to test the distributed memory models’ prediction that performance on the old/new test format with targets and similar lures would be the most similar to the A-B’ test format. Accordingly, we ran two separate regressions between performance on the old/new test format with targets and similar lures vs. the A-A’ and A-B’ test formats (see Equations (24) and (25)). We then compared which of the two slopes was closer to 1. We next compared the raw performance values between the test formats and we performed contrasts between the conditions to determine which conditions were the most similar. Additionally, we used a dendrogram analysis in which we calculated the Euclidean distance between performance on the various test formats. In accordance with the distributed memory models, we predicted that these analyses would reveal that the A-B’ and the old/new test format with targets vs. similar lures would be the most similar, followed by the A-A’ test format. Likewise, we predicted that the A-X and the old/new with targets and novel foils would be similar to each other.

We tested the prediction of the distributed memory models that performance would be correlated across all of the forced-choice and the old/new test formats. Specifically, although previous research suggests that there are dissociations in performance on these test formats, the distributed memory models predict that performance will be correlated between all test formats (i.e., as a function of the learning rate of the model/participant). Therefore, positive correlations between the test formats would support the predictions of the distributed memory models while null correlations would support the notion that these test formats recruit distinct cognitive and neural processes.

### Results and Discussion

We designed two versions of the Mnemonic Similarity Task (Stark et al., 2013, 2015, 2019): the old/new MST and the forced-choice MST (see Figure 1). In both tasks, participants first completed an incidental encoding phase in which they made indoor/outdoor judgments for each image. During the test phase of the old/new MST, participants saw repeated items (targets), similar items (similar lures), and novel items (novel foils), and we asked them to respond “old” for items that were exactly the same as one they saw during the indoor/outdoor task and to respond “new” to items that were similar to or totally new from any images they saw during the indoor/outdoor task. We used a within-participants design, in which each participant completed both versions of the MST in a randomized order (N=145). We report the results of each version as well as the comparisons between the tasks below.

#### Forced-Choice Data

We first investigated whether there was an effect of test format on performance of the forced-choice version of MST (see Figure 1b) using a generalized linear mixed model with a binomial family. Specifically, we first compared a null model (random slopes and intercepts) with a test-format model that incorporated the mean performance from Experiment 2 in Huffman and Stark (2017): A-X: logit(0.959) A-A’: logit(0.836), A-B’: logit(0.756). A likelihood ratio test revealed that the test-format model accounted for significantly more variance than the intercept-only model (*χ*^2^(2*, N* = 145) = 193.69, *p <* 2.2 × 10*^−^*^16^; intercept-only model *BIC* = 2, 284.2, test-format model *BIC* = 2, 096.6; see Figure 12a). The fixed-effects beta estimates for the test-format model were an intercept of -0.186 and a slope of 1.09. Thus, these results replicate our previous findings and suggest that the magnitude of our observed effects here are similar (i.e., our intercept is close to zero and our slope is close to 1). We next compared our test-format model with the predictions of the CLS hippocampal model of equal performance on the A-A’ and A-B’ test (A-A’ = A-B’: mean(logit((0.836, 0.756))); see Figure 2a; cf. Norman & O’Reilly, 2003; Migo et al., 2009). We compared the Bayesian information criterion (BIC; lower values indicate model preference) and found evidence that was very much in favor of the distributed memory models vs. the CLS hippocampal model (BIC test-format-model: 2,096.6; BIC CLS hippocampal model: 2,222.1; *BIC_distributed_* − *BIC_hippocampus_* = −125.5). Moreover, we used the BIC approximation for calculating Bayes factors (see Equation (2); Wagenmakers, 2007; Jarosz & Wiley, 2014) and we found that the evidence was in favor of the distributed model vs. the hippocampal model (*BF*_10_ ≈ 1.74 × 10^27^; where 1 indicates distributed memory models and 0 indicates the CLS hippocampal model), thus suggesting that the data are 1.74 × 10^27^ times more likely for the predictions of the distributed memory models relative to the predictions of the CLS hippocampal model.

We next performed planned follow-up tests to determine whether performance on the A-X format was better than the A-A’ format and whether performance on the A-A’ format was better than the A-B’ format (i.e., as in Huffman & Stark, 2017). First, we refit the null model and the test-format model while only including data from the A-X and A-A’ test formats. We again found that the test-format model accounted for significantly more variance than the intercept-only model (*χ*^2^(1*, N* = 145) = 173.15, *p <* 2.2 × 10*^−^*^16^; null model *BIC* = 1, 516.3, test-format model *BIC* = 1, 348.8; intercept = 0.052, slope = 0.98), thus indicating that performance on the A-X test format is significantly better than performance on the A-A’ test format. Second, we refit the intercept-only model and the test-format model while only including data from the A-A’ and A-B’ test format. We again found that the test-format model accounted for significantly more variance than the intercept-only model (*χ*^2^(1*, N* = 145) = 97.49, *p <* 2.2 × 10*^−^*^16^; null model *BIC* = 1, 660.2, test-format model *BIC* = 1, 568.4; intercept = -0.45, slope = 1.29), thus indicating that performance on the A-A’ test format is significantly better than performance on the A-B’ test format. Altogether, these results replicate our previous findings (Huffman & Stark, 2017), and strongly suggest that there is an effect of test format, thus further bolstering the notion that participants perform better on the A-A’ test format than the A-B’ test format. Therefore, these results are consistent with the predictions of the distributed memory models (MINERVA 2: Figure 5b, TODAM: Figure 6b, both versions of the Hopfield/autoassociative neural network: Figure 7b and Figure 8b; also see the neocortical model from Simulation 4 in Norman & O’Reilly, 2003) but not with the CLS hippocampal model (see Figure 2a; Norman & O’Reilly, 2003).

We next investigated the effect of target-lure similarity, determined by lure bin levels, on the performance (see Figure 1c). We focus here on the results of A-A’ and A-B’ test formats because the A-X test format trials did not manipulate the similarity between the encoding objects and test objects. For this analysis, we determined target-lure similarity using the lure bin levels, which were generated from the empirical mnemonic similarity from previous experiments (i.e., defined by the p(“old”) to each stimulus, which is a measure of confusability; the lure bins were generated from the old/similar/new test format: Lacy et al., 2011; Stark et al., 2013, 2015, 2019). We compared the test-format model with a test-and-lure-bin model and found that the test-format-and-lure-bin model accounted for significantly more variance than the test-format model (*χ*^2^(1*, N* = 145) = 135.24, *p <* 2.2 × 10*^−^*^16^; test-format model *BIC* = 4, 410, test-format-and-lure-bin model *BIC* = 4, 282), thus demonstrating that participants’ performance varied as a function of the lure bins (i.e., target-lure similarity; see Figure 12b). We next tested whether the effect of the lure bin varied as a function of test format (e.g., was more pronounced in one test format). Interestingly, a test-format by lurebin interaction model did not account for significantly more variance than the test-and-lure-bin model (*χ*^2^(1*, N* = 145) = 0.13, *p* = 0.72; test-format by lure-bin interaction model *BIC* = 4, 289.2, note that the BIC for this model is higher than the test-format-and-lure-bin model), thus suggesting that the lure bin effect was similar between the A-A’ and A-B’ test formats (*BF*_01_ ≈ 36.6). To further explore the similarity of the lure bin effect on the A-A’ and A-B’ test formats, we next compared a lure bin model with an intercept-only model within each test format separately. We found that the lure-bin model accounted for significantly more variance than the intercept-only model for both the A-A’ test format (*χ*^2^(1*, N* = 145) = 88.71, *p <* 2.2 × 10*^−^*^16^; intercept = 1.70, lure bin beta = 0.26; intercept-only model *BIC* = 2, 080.7, lure-bin model *BIC* = 1, 998.5) and the A-B’ test format (*χ*^2^(1*, N* = 145) = 126.71, *p <* 2.2 × 10*^−^*^16^; intercept = 1.05, lure bin beta = 0.26; intercept-only *BIC* = 2, 434.9, lure-bin model *BIC* = 2, 310.2). Note that the beta estimates for the lure bin effect were remarkably similar across these two test formats: 0.26 for both the A-A’ and the A-B’ test formats. Taken together, these results clearly indicate that there is an effect of the lure bin on performance of both the A-A’ and the A-B’ test format and further suggest that this effect is similar across these two formats.

#### Old/New Data

We next investigated whether there was an effect of test format on performance on the old/new version of the MST (for the raw proportion of responses on each trial, see Figure 13a). We calculated the area under the ROC curve, *A_z_* (see *Comparison of the Forced-Choice and Old/New Test Formats*) and then applied the logit transformation to the resultant values to compare performance between conditions. We found that performance was significantly better than chance for the target/novel foil discrimination (mean= 2.527, *t*_144_ = 25.96, *p <* 2.2 × 10*^−^*^16^; *BF*_10_ ≈ 4.54 × 10^52^ [the t was very large, so the function used an approximation]; Cohen’s d= 2.16; see the *A_z_* old/new in Figure 16) and for the target/similar lure discrimination (mean= 1.20, *t*_144_ = 23.20, *p <* 2.2×10*^−^*^16^; *BF*_10_ ≈ 1.08 × 10^47^ [the t was very large, so the function used an approximation], Cohen’s d= 1.93; see the *A_z_*old/lure in Figure 14b). We also found that performance was significantly better for the target/novel foil discrimination than the target/similar lure discrimination (mean difference= 1.33, *t*_144_ = 20.92, *p <* 2.2 × 10*^−^*^16^, *BF*_10_ ≈ 1.22 × 10^42^ [the t was very large, so the function used an approximation], Cohen’s d= 1.737; also see *Comparison of the Forced-Choice and Old/New Test Formats* for the comparison of the forced-choice and old/new performance).

**Figure 16:**
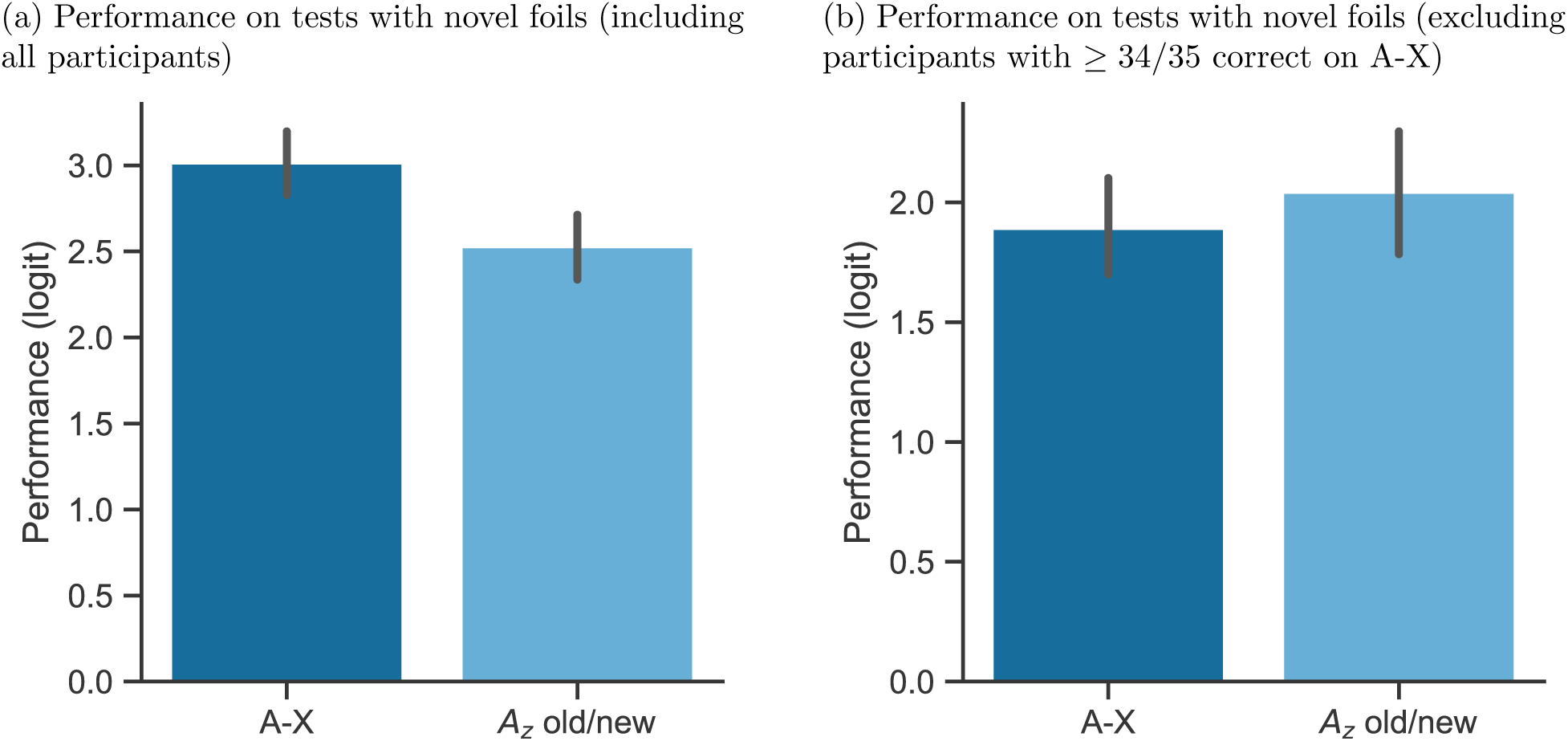
Comparison of human performance on the A-X and old/new test format with targets and novel foils indicated that performance was better on the A-X test format, but this might be driven by a ceiling effect in the A-X test format. (a) When we included all participants, we found that A-X performance was significantly better than *A_z_* with targets and novel foils. (b) However, when we tested whether this could be driven by a ceiling effect (here, we limited our analysis to participants with fewer than 34/35 correct on the A-X format), we observed that the A-X *> A_z_* old/new advantage failed to reach significance, thus suggesting a ceiling effect was driving the original effect. Overall, these results are consistent with the predictions of the distributed memory models (see Figure 9 and Figure 10). Bars represent the mean and the error bars indicate the 95% confidence intervals.

We next tested for our expected effect of a relationship between target-lure similarity and performance on the A-A’ and A-B’ test formats. Here, we fit a generalized linear model (binomial family) to the proportion of “new” responses to similar lures (note: the correct answer is “new”, while the incorrect response is “old” because these are all similar lures; see Figure 1c). We found that the lure-bin model (based on the empirically defined mnemonic similarity; Lacy et al., 2011; Stark et al., 2013, 2015, 2019) accounted for significantly more variance than the intercept-only model for the similar lure data (*χ*^2^(1*, N* = 145) = 195.93, *p <* 2.2 × 10*^−^*^16^; intercept = 0.31, *β_lure bin_* = 0.41; null model *BIC* = 3, 353.4, lure-bin model *BIC* = 3, 141.2; see Figure 13b). Therefore, these results replicate and extend previous work that showed that performance varies in a monotonic manner with the differences of lure bins (i.e., mnemonic similarity) on the old/similar/new test format as well as our findings here on the A-A’ and A-B’ forced-choice test formats. Altogether, our results of the artificial neural network suggest that the transfer function between memory performance and the stimulated target-lure pattern similarity in area IT is approximately linear (see Quantification of Target/Lure Similarity with an Artificial Neural Network and Figure 4), which extends to the A-A’, A-B’, and old/new test format with targets and similar lures.

#### Comparison of the Forced-Choice and Old/New Test Formats

We next aimed to compare performance between the forced-choice and old/new test formats to test the predictions from the distributed memory models vs. the CLS hippocampal model. The distributed memory models predict that performance on the old/new test format with targets and similar lures will be most similar to the A-B’ test format. Specifically, in the case of running many simulations and calculating the average performance at the group level (i.e., akin to a fixed-effects analysis), the distributed memory models predict that performance on the old/new test format with targets and similar lures will be equivalent to the A-B’ test format, which is in line with previous theoretical and empirical findings for traditional recognition memory paradigms with targets and novel foils (Green & Moses, 1966; Swets & Pickett, 1982; Stanislaw & Todorov, 1999). Note that in contrast to the predictions of the distributed memory models, the CLS hippocampal model predicts equivalent performance on the old/new test format with targets and similar lures, the A-A’ test format, and the A-B’ test format. Moreover, the distributed memory models predict that performance will be correlated across all of the test formats, including the formats with targets vs. novel foils (i.e., driven by individual differences in the learning rate). Thus, we performed several analyses to test the similarity between the forced-choice and old/new test formats.

First, we tested the distributed memory models’ prediction that performance on the old/new test format with targets and similar lures would be closest to the A-B’ test format. We first calculated the area under the receiver operating characteristic (ROC) curve, which is referred to as *A_z_* (see *Comparison of the Forced-Choice and Old/New Test Formats*). We then converted the performance scores with the logit function. We then ran a two-stage regression model to estimate the slope and intercepts of the relationship between the test formats with targets and similar lures. We observed that the estimated slope was relatively close to 1 for the comparison between the *A_z_* for the old/similar lures and the A-B’ test format (estimated slope= 1.49, estimated intercept= −0.71), which is in line with the predictions of the distributed memory models. Likewise, the estimated slope for the comparison between the *A_z_* for the old/similar lures and the A-A’ test format was (numerically) larger (estimated slope= 1.98, estimated intercept= −0.61), which is also in line with the predictions of the distributed memory models (see Figure 14a). To further assess the similarity between *A_z_* for old/similar lures and the A-B’ test formats, we next performed difference tests to compare the formats with targets and similar lures. A paired t-test revealed that performance was significantly better on the A-A’ test format than the *A_z_* for targets and similar lures (mean difference= 0.558, *t*_144_ = 7.87, *p* = 7.91 × 10*^−^*^13^; *BF*_10_ = 9.16 × 10^9^; Cohen’s d= 0.653). A paired t-test also revealed that the *A_z_* for targets and similar lures was (numerically) greater than performance on the A-B’ test format (mean difference= −0.129, *t*_144_ = −2.24, *p* = 0.026) but the magnitude of this effect was weak overall (*BF*_10_ = 1.04, Cohen’s d= 0.186; see Figure 14b). To further test whether performance of the *A_z_* for targets and similar lures was closer to the A-B’ test format than the A-A’ test format, we next performed a difference test. We found that the difference between the A-A’ vs. *A_z_* for targets and similar lures was significantly greater than the difference between the *A_z_* for targets and similar lures vs. the A-B’ test format (mean difference= 0.429, *t*_144_ = 3.92, *p* = 0.00014, *BF*_10_ = 119.1, Cohen’s d= 0.325; see Figure 14c). Altogether, these results suggest that performance on the *A_z_* for targets and similar lures is numerically closer to the A-B’ test format than the A-A’ test format, thus supporting the predictions of the distributed memory models.

We next went back to the computational models to determine whether the empirical effect of slightly enhanced performance on the old/new test format with targets and similar lures vs. the A-B’ test format could be explained within the distributed memory models. Briefly, in our original Performance on tests with similar lures simulations, we combined the data from all of the simulated participants in our calculations of performance between the test formats. Under those conditions, which are akin to a fixed-effects analysis, we found that performance on the old/new test format with targets and similar lures was equivalent to performance on the A-B’ test format. Additionally, this is a case with a (very) large number of data points for each condition. Thus, we were interested in modeling the effects at the level of individual participants (i.e., akin to a random-effects analysis). Interestingly, we found that the models predicted a significant advantage for the old/new test format with targets and similar lures relative to the A-B’ test format in this analysis. Specifically, the models predicted that performance on the A-A’ test format is better than the *A_z_* with targets and similar lures, the *A_z_* with targets and similar lures is better than A-B’, and the difference between the A-A’ and *A_z_* is great than the difference between *A_z_* and A-B’ (for example, see the results from MINERVA 2 in Figure 15; the results were the same in the other distributed memory models). Therefore, while this was not part of our pre-experimental modeling (and thus we temper our interpretations here), we think that these results provide further support for the notion that the distributed memory models can capture performance on various versions of the Mnemonic Similarity Task.

We next aimed to further examine the relationship between performance on all five test formats. (a) Performance on tests with novel foils (including all participants)

A dendrogram analysis of the Euclidean distance between performance on all 5 test formats revealed that *A_z_* for targets and similar lures was most similar to the A-B’ test format (see Figure 14d) and the A-A’ test format was in a similar branch of the dendrogram results as the *A_z_* for targets and similar lures as well as the A-B’ test format. Likewise, the *A_z_* for targets and novel foils was in cluster with the A-X test format. For completeness, we compared the *A_z_* for targets and novel foils vs. the A-X test format and we found that performance was significantly better on the A-X test format (mean difference= 0.487, *t*_144_ = 4.80, *p* = 3.91 × 10*^−^*^6^, *BF*_10_ = 3, 356.1, Cohen’s d= 0.399; see Figure 16a). However, we wondered whether this could be a spurious ceiling effect driving this difference in the A-X test format. Specifically, a large number of participants (81 of 145) exhibited perfect (35/35) or near perfect (34/35) performance on the A-X test format. Therefore, we tested whether there was still a test format effect in participants that were correct on fewer than 34 of the 35 trials on the A-X test format (which resulted in 64 participants in the analysis and 81 excluded participants). Interestingly, the apparent A-X *> A_z_* old/new effect disappeared under these conditions (mean difference= −0.151, *t*_63_ = 1.22, *BF*_01_ = 3.60 [i.e., in favor of the null hypothesis], Cohen’s d= 0.152; see Figure 16b), thus suggesting that a ceiling effect was driving the difference between these test formats. Altogether, the finding of clustering in the dendrogram analysis matches the predictions of the distributed memory models.

We next aimed to test the prediction from the distributed memory models that performance would be correlated on all of the forced-choice and old/new test formats. Consistent with the predictions of the distributed memory models, we found a significant correlation between all test formats (see Figure 17). Specifically, all of the forced-choice test formats exhibited a significant correlation (A-X vs. A-A’: *r* = 0.62, *t*_143_ = 9.37, *p <* 2.2 × 10*^−^*^16^, *BF*_10_ = 3.61 × 10^13^; A-X vs. A-B’: *r* = 0.46, *t*_143_ = 6.13, *p <* 8.27 × 10*^−^*^9^, *BF*_10_ = 1.70 × 10^6^; A-A’ vs. A-B’: *r* = 0.53, *t*_143_ = 7.48, *p <* 6.69 × 10*^−^*^12^, *BF*_10_ = 1.38 × 10^9^). Moreover, all three forced-choice test formats exhibited a significant correlation with the old/new test format with targets and novel foils (A-X vs. *A_z_* old/new: *r* = 0.46, *t*_143_ = 6.26, *p <* 4.25 × 10*^−^*^9^, *BF*_10_ = 3.18 × 10^6^; A-A’ vs. *A_z_* old/new: *r* = 0.41, *t*_143_ = 5.41, *p <* 2.56 × 10*^−^*^7^, *BF*_10_ = 6.91 × 10^4^; A-B’ vs. *A_z_* old/new: *r* = 0.36, *t*_143_ = 4.63, *p <* 7.98 × 10*^−^*^6^, *BF*_10_ = 2.86 × 10^3^) and the old/new test format with targets and similar lures (A-X vs. *A_z_*old/sim: *r* = 0.42, *t*_143_ = 5.51, *p <* 1.60 × 10*^−^*^7^, *BF*_10_ = 1.07 × 10^5^; A-A’ vs. *A_z_* old/new: *r* = 0.47, *t*_143_ = 6.43, *p <* 1.77 × 10*^−^*^9^, *BF*_10_ = 7.24 × 10^6^; A-B’ vs. *A_z_* old/new: *r* = 0.42, *t*_143_ = 5.48, *p <* 1.86 × 10*^−^*^7^, *BF*_10_ = 9.32 × 10^6^). Finally, the comparison between the two old/new formats relies upon data that are not independent, but we report the correlation here for completeness (*r* = 0.81). Altogether, our finding of significant correlations between all of the test formats supports the predictions of the distributed memory models.

**Figure 17:**
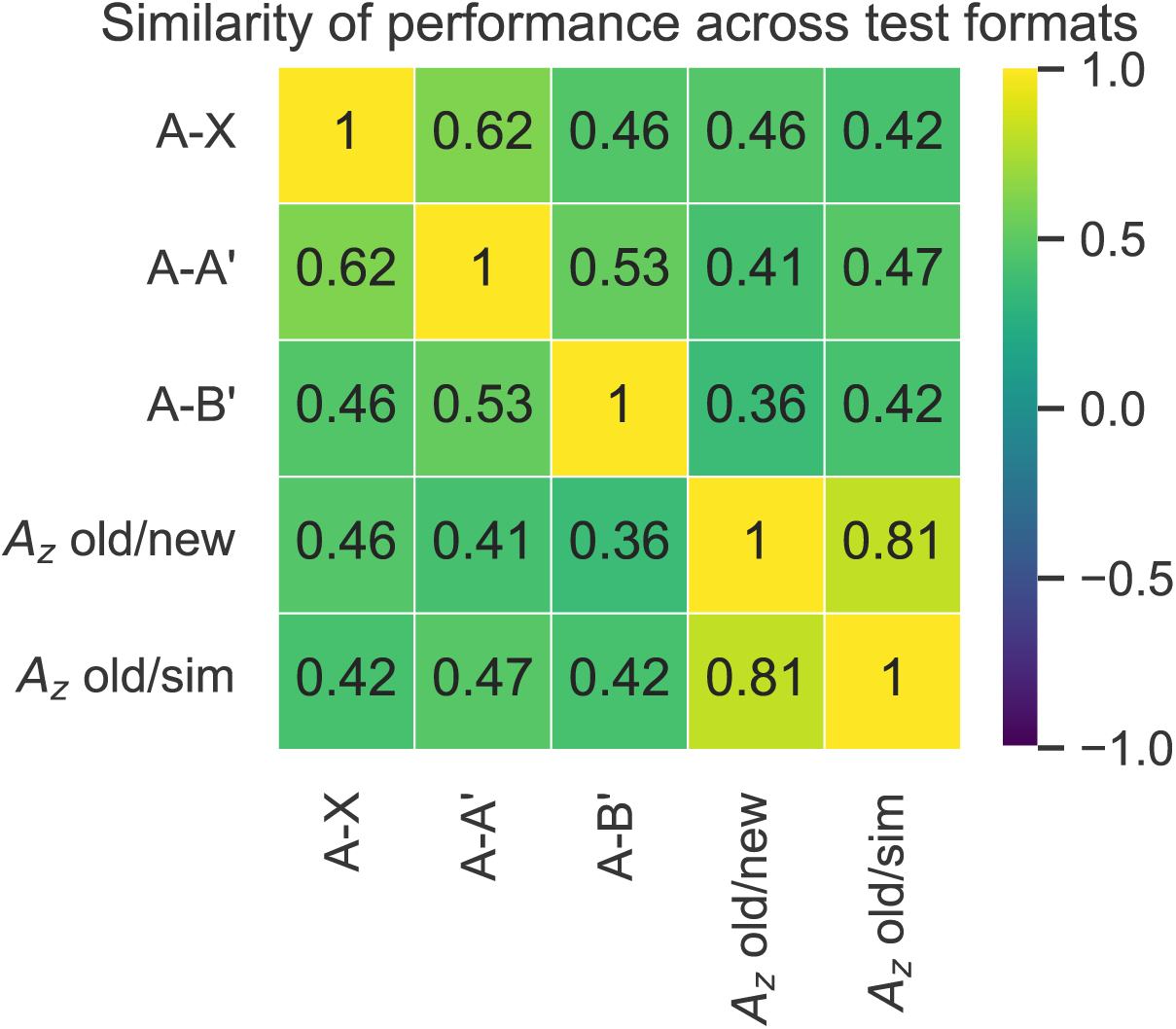
We observed significant correlations between human performance on all test formats, thus supporting the predictions of the distributed memory models (i.e., compare the results here to Figure 11; note that the comparison of the *A_z_* old/new and *A_z_* old/sim are from data that are not independent).

## General Discussion

As we discussed above, there is near consensus that the brain contains different systems that play different roles in memory: a hippocampal system that employs pattern separation and is geared toward avoiding interference between similar experiences and a neocortical system that employs distributed representations and is geared toward allowing generalization between similar experiences; however, it was hitherto unknown whether memory-based decisions are primarily driven by the pattern-separated representations of the hippocampus (e.g., dentate gyrus) vs. distributed representations (e.g., neocortex, CA1, CA3) vs. a composite representation between these systems. Therefore, we aimed to develop and test the predictions between competing computational models of human performance on memory tests with targets, similar lures, and novel foils. Moreover, we believe that these test formats provide a critical testbed for comparing the representations that support human memory. Specifically, we compared predictions of the CLS hippocampal model (Norman & O’Reilly, 2003; Norman, 2010)—which emphasizes sparse, pattern-separated representations for avoiding interference between similar memories—vs. distributed memory models—which emphasize distributed representations (see our modeling results above). We focused our empirical analyses on areas in which the two classes of models make competing predictions: 1) the nature of the transfer function between performance and target-lure similarity, 2) whether or not there would be a significant effect of test format on performance of memory tests with targets and similar lures, and 3) whether different memory tasks would produce correlated or dissociable performance via individual differences in performance. Our results clearly favored the predictions of the distributed memory models over the predictions of the CLS hippocampal model: 1) we found an approximately linear transfer function between performance and the simulated target-lure similarity in an artificial neural network of area IT, 2) we observed systematic differences in performance between the forced-choice and old/new test formats, and 3) individual differences analyses suggest that differences in learning rate lead to correlated performance on all 5 test formats, which we discuss in turn below. Therefore, our results suggest that human memory performance is primarily driven by distributed representations (e.g., of the neocortex or subregions other than the dentate gyrus of the hippocampus [e.g., CA3, CA1]). We also discuss implications of our modeling and empirical results for future research using behavioral, neural, and computational modeling approaches for studying interference resolution in human memory.

### The Transfer Function between Memory Performance and the Simulated Target-Lure Pattern Similarity in Area IT Appears to be Approximately Linear

Tasks that include targets and similar lures typically aim to tap into the pattern-separation mechanisms of the hippocampus (e.g., the Mnemonic Similarity Task [MST]: Kirwan & Stark, 2007; Stark et al., 2013, 2015, 2019; the MST had been used in more than 100 behavioral and neuroimaging publications as of 2019: Stark et al., 2019). The CLS hippocampal model and the distributed memory models make competing predictions about the degree to which target-lure similarity will influence memory performance. Specifically, the CLS hippocampal model predicts that humans will only exhibit interference when target-lure similarity is very high or when the similarity between all items in a list is very high (see Figure 2a and Norman & O’Reilly, 2003; Norman, 2010). We aimed to generate an objective measure of the target-lure similarity using a biologically inspired artificial neural network, CORnet-S (Kubilius et al., 2019; Muttenthaler & Hebart, 2021), to compare the pattern similarity of simulated activity within area IT, which is heavily interconnected with perirhinal cortex (Suzuki & Amaral, 1994). Specifically, extent models of the medial temporal lobe predict that area IT would project strongly into the perirhinal cortex (Suzuki & Amaral, 1994), which would then project to entorhinal cortex and into the hippocampus (e.g., Eichenbaum et al., 2007; Ranganath & Ritchey, 2012; Squire, Stark, & Clark, 2004; Buffalo, Bellgowan, & Martin, 2006; Rolls, 2018). Models of hippocampal pattern separation posit that the hippocampus, and the dentate gyrus in particular, will decrease the pattern similarity between targets and similar lures relative to its input patterns from the neocortex (e.g., McClelland et al., 1995; O’Reilly & Norman, 2002; Norman & O’Reilly, 2003; Norman, 2010; O’Reilly & McClelland, 1994). Therefore, if human participants rely on the pattern-separated representations of the dentate gyrus to make memory judgments (i.e., based on previous neuroscientific findings; e.g., J. K. Leutgeb et al., 2007; Neunuebel & Knierim, 2014; S. Leutgeb & Leutgeb, 2007; Lacy et al., 2011; Yassa & Stark, 2011; Reagh et al., 2018), then we would expect behavior to follow a nonlinear function and only exhibit interference for very high levels of target-lure similarity in its input regions (see Figure 2a and Norman & O’Reilly, 2003; Norman, 2010). Conversely, as we reported here, the distributed memory models predict that the degree of target-lure similarity will exhibit a strong, monotonic, and relatively continuous influence on memory performance (MINERVA 2: Figure 5b, TODAM: Figure 6b, the familiarity-based Hopfield/autoassociative neural network: Figure 7b, and the recall-based Hopfield/autoassociative neural network: Figure 8b; also see the medial temporal lobe cortex model from Norman & O’Reilly, 2003; Norman, 2010). Therefore, testing these competing predictions can shed important light on the underlying mechanisms of human memory.

Our results provide evidence for what appears to be a linear mapping between mnemonic discrimination and simulated target-lure pattern similarity in area IT. Specifically, when we compared the resultant simulated pattern similarity in area IT to the original behavioral data that was used to generate the lure bins of the stimuli in the Mnemonic Similarity Task (Kirwan & Stark, 2007; Stark et al., 2013, 2015, 2019), we observed evidence of an approximately linear transfer function between mnemonic discrimination performance and the simulated target-lure similarity in area IT (see Figure 4). Moreover, in our current empirical data, we replicated the effect of the influence of lure bins on memory performance on the old/new test format with targets and similar lures (see Figure 13b; e.g., Lacy et al., 2011; Stark et al., 2013, 2015, 2019). Here, we also extended the finding of an effect of the lure bin on performance of both the A-A’ and A-B’ test formats (see Figure 12b). The finding that the lure similarity effect was not significantly different between the A-A’ and A-B’ test formats (and that it was significant in both test formats independently) suggests that target-lure similarity affects performance across all of these test formats. Moreover, combined with the simulated pattern similarity results in area IT, our results further suggest that the transfer function between mnemonic discrimination and target-lure similarity is approximately linear in the old/new, A-A’, and A-B’ test formats.

While our study is the first (to our knowledge) to test the transfer function between mnemonic discrimination performance and target-lure similarity using an artificial neural network, Motley and Kirwan (2012) created a parametric version of the Mnemonic Similarity Task in which they modulated the target-lure similarity by parametrically rotating the objects to form the similar lures (rotations: 15*^◦^*, 25*^◦^*, 35*^◦^*, and 55*^◦^*). Interestingly, they found that behavioral performance (the proportion of times that participants responded “old” as well as “rotated”=“similar”) varied linearly as a function of the target-lure rotation. Therefore, our results extend these findings from their parametric manipulation of target-lure similarity. Given the widespread use of the Mnemonic Similarity Task and other similar measures, we argue that our results provide important insight into the nature of the underlying representations that support performance on the task. Specifically, our results suggest that mnemonic discrimination performance is primarily driven by distributed representations (e.g., the neocortex, CA3, CA1) rather than the pattern separation mechanisms of the dentate gyrus or a composite representation across these systems.

As we discussed in the Introduction, previous fMRI research using the Mnemonic Similarity Task (and related paradigms) have generally found a nonlinear mapping between the response profile of the dentate gyrus/CA3 and the mnemonic similarity of target-lure pairs but a linear relationship between the response profile of CA1 and the mnemonic similarity of target-lure pairs (e.g., Lacy et al., 2011; Reagh et al., 2018; for review see: Yassa & Stark, 2011). Therefore, given that mnemonic similarity was operationally defined as mnemonic discrimination (i.e., 1-p(‘old’); Lacy et al., 2011; Stark et al., 2013, 2015, 2019), these findings also reveal a potential disconnect between the dentate gyrus/CA3 and memory performance. In contrast, the linear relationship between CA1 activity and mnemonic similarity suggests that CA1 may play a stronger role in behavioral performance. Moreover, previous computational modeling and empirical research suggests that CA1 might play a role in match-mismatch detection (e.g., via a match-mismatch mechanism; Hasselmo et al., 1996; Meeter et al., 2004) and previous research has shown a linear transfer function between response profiles of CA1 and environmental/stimulus similarity (e.g., Guzowski et al., 2004; Kumaran & Maguire, 2006, 2007; Duncan et al., 2012). Therefore, a possible explanation for our findings here is that participants rely on neural signals other than pattern-separated representations in the dentate gyrus to make memory-based judgments on tasks with targets and similar lures. For example, the underlying representations of the distributed models that we use here are similar to the representations of the neocortex (e.g., perirhinal cortex e.g., see the neocortical model from O’Reilly & Norman, 2002; Norman & O’Reilly, 2003; Norman, 2010) and CA1. Additionally, given the functional similarity of the Hopfield/autoassociative neural network to area CA3 (cf. Rolls, 2007; Marr, 1971; McNaughton & Morris, 1987; Wilson et al., 2006), it is possible that CA3 might play a key role in memory performance on these tasks. Moreover, a recall-based version of the Hopfield/autoassociative neural network was previously shown to account for performance on recall tasks (i.e., in addition to the recognition memory tasks that we used here; e.g., Rizzuto & Kahana, 2001), thus suggesting it could potentially be a more general memory mechanism. Finally, it is possible that memory-based decisions are driven by the accumulation of evidence throughout the brain (i.e., in a more holistic, global manner; cf. Squire et al., 2007; Wixted, 2007; Norman & O’Reilly, 2003; Norman, 2010).

While the CLS model predicts that the hippocampus will generally not be useful for computing the global similarity between stimuli (i.e., given its use of pattern separation; Norman & O’Reilly, 2003; Norman, 2010), as we discussed in the Introduction, Davis et al. (2014) found that the global similarity between targets and foils in the hippocampus and other areas of the medial temporal lobe was related to memory-based decisions in a recognition memory paradigm. Therefore, their results suggest that there are cases in which the hippocampus would be able to compute the global match, which would, in turn, be related to memory-based decisions. Combined with our findings here, we argue that it will be interesting for future research to further investigate the degree to which we make memory decisions based on the pattern-separated representations in the dentate gyrus vs. distributed representations in the neocortex (or the hippocampus) vs. combined evidence from all of these sources (cf. Squire et al., 2007; Wixted, 2007; Norman & O’Reilly, 2003; Norman, 2010).

In summary, the results of the transfer function between mnemonic discrimination performance and target-lure similarity in input regions of the hippocampus appears to be linear, which is better fit by the predictions of the distributed memory models than by the pattern-separation mechanisms of the dentate gyrus. Moreover, we argue that when we reexamine the findings from previous fMRI studies, the evidence actually points toward a closer relationship between mnemonic discrimination performance and activity in areas outside of the dentate gyrus (e.g., CA1, perirhinal cortex). We return to the discussion of other possible reasons that the behavior did not fit the predictions of the standard version of the CLS hippocampal model (see The Test Format Effect on the Forced-Choice Tests Suggests that Memory Performance is Primarily Driven by Distributed Representations and Avenues for Testing the Generality of our Empirical and Modeling Results).

### The Test Format Effect on the Forced-Choice Tests Suggests that Memory Performance is Primarily Driven by Distributed Representations

We compared the performance of several computational models at accounting for human behavioral performance on different test formats of the Mnemonic Similarity Task (the CLS hippocampal model: Figure 2a; MINERVA 2: Figure 5b, TODAM: Figure 6b, the familiarity-based Hopfield/autoassociative neural network: Figure 7b, and the recall-based Hopfield/autoassociative neural network: Figure 8b; also see the medial temporal lobe cortex model from Norman & O’Reilly, 2003; Norman, 2010). One of our key findings is an effect of differences in performance as a function of the test format. For example, for the forced-choice test formats, performance was best on the A-X test format, followed by the A-A’ test format, followed by the A-B’ test format (see Figure 12a). All of the models that can account for this effect of test format—and in particular the A-A’ *>* A-B’ effect—suggest that encoding variability plays a key role in this effect (cf. Simulation 4 from Norman & O’Reilly, 2003; Hintzman, 1988; Huffman & Stark, 2017). Specifically, the models predict that encoding variability leads to an increased number of errors on the A-B’ test format because sometimes the encoding of the original B item exceeds the encoding of the original A items. In this situation, the response to the B’ item can trigger a stronger global memory signal because it matches more features in memory. For example, when encoding variability is minimized, the difference between the A-A’ and A-B’ test formats diminishes (cf. Figure 4 from Huffman & Stark, 2017 with MINERVA 2). Interestingly, Rollins et al. (2019) used eye tracking to test whether performance on the A-B’ test format varies as a function of encoding variability, as indexed by the number of fixations. Specifically, they found a significant test-format (A-A’ vs. A-B’) by subsequent memory accuracy (correct vs. incorrect) interaction. A follow-up t-test showed that participants made significantly more fixations (a measure of encoding strength) to B items than A items on subsequently incorrect trials of the A-B’ forced-choice test format. Therefore, their findings supported a key prediction from the distributed memory models that the impaired performance on the A-B’ test format relative to the A-A’ test format is influenced by encoding variability (i.e., better encoding of the original B item than the original A item). Future research can further test these predictions using other measures of encoding strength (e.g., subsequent memory effects with fMRI and EEG). Counter to the prediction of an effect of encoding variability leading to better performance on the A-A’ test format relative to the A-B’ test format, the standard CLS hippocampal model predicts that performance will be similar on these two test formats (see Figure 2a; Norman & O’Reilly, 2003; Migo et al., 2009) even with simulated encoding variability (see Simulation 4 from Norman & O’Reilly, 2003). In fact, the CLS model predicts that encoding variability will influence performance in the neocortical model but not the hippocampal model. They suggest that the pattern separation and recall-to-reject mechanisms of the hippocampus would give it two opportunities to reject the items in the A-B’ test condition (i.e., once for the A item and once for the B’ item), thus any benefit of the covariance of the items in the A-A’ test format is offset by the additional ability for recall-to-reject on A-B’ test format.

Although our behavioral results here as well as in other studies (Hintzman, 1988; Jeneson et al., 2010; Tulving, 1981; Huffman & Stark, 2017; Rollins et al., 2019; Fandakova et al., 2021) are discordant with the CLS hippocampal model’s prediction of a null effect of a difference between performance on the A-A’ vs. A-B’ test formats, we suggest that there could be modifications under which the CLS hippocampal model would account for the extant behavioral findings. First, the output of the hippocampal model and the neocortical model could be combined and a separate neural network could be used to model decision making (cf. Norman & O’Reilly, 2003; Norman, 2010). In this situation, the combined output could predict an A-A’ *>* A-B’ test format because the neocortical model already predicts such an effect (i.e., when there is encoding variability). As a second approach, we predict that different learning rules for implementing encoding variability might lead to the A-A’ *>* A-B’ test format effect. Norman and O’Reilly (2003) implemented encoding variability by universally scaling the learning-rate parameter on all of the weights. Specifically, for each trial, they multiplied the learning rate by a variable encoding strength (see Simulation 4 from Norman & O’Reilly, 2003). In effect, this changes the encoding strength of all features uniformly. In contrast, other approaches for implementing encoding variability could be to use either a connection-wise probabilistic encoding rule (e.g., as applied here for the Hopfield/autoassociative neural network; Rizzuto & Kahana, 2001) or a feature-wise probabilistic encoding rule (e.g., as applied here for MINERVA 2 and TODAM). Specifically, the connection-wise probabilistic learning rule would essentially model random variability in terms of how many connections are modified. Likewise, the feature-wise probabilistic learning rule would essentially model random variability in terms of whether or not stimulus features were properly activated or stored during the original encoding phase (e.g., we can imagine that we neither pay attention to nor memorize every feature in any given trial). In either situation, the number of features as well as which specific features are encoded would be variable on each trial. For example, the Hopfield/autoassociative neural network that we used here predicted a test format effect. In effect, the connection-wise or the feature-wise probabilistic learning rule would also allow the encoding variability to interact with the sampling variability (e.g., in some cases encoding more of the inactive units than the active units). Interestingly, encoding variability within Hopfield/autoassociative neural networks was also found to be important for associative recall tasks, thus suggesting a possible extension and connection to different behavioral paradigms (Rizzuto & Kahana, 2001). Given the structural similarity of the Hopfield/autoassociative neural network to the CA3 subregion of the hippocampus (cf. Rolls, 2007; Marr, 1971; McNaughton & Morris, 1987; Wilson et al., 2006), we think it is possible that such a change to the hippocampal network will account for the test-format effects that we observed in our empirical data, and we plan to explore this possibility in future work.

### The Relationships between Performance on the Forced-Choice and Old/New Test Formats Suggests that Memory Performance is Primarily Driven by Distributed Representations

The distributed memory models predict specific relationships between performance on the forced-choice and old/new test formats: performance on the old/new test format with targets and novel foils will be similar to performance on the A-X test format, and performance on the old/new test format with targets and similar lures will be significantly worse than performance on the A-A’ test format and most similar to performance on the A-B’ test format (see Figure 10 and Figure 15). Conversely, the standard CLS hippocampal model predicts that performance will be virtually the same across all of the test formats with targets and similar lures. Our results fit with the predictions of the distributed memory models (see Figure 14 and Figure 16). Moreover, the distributed memory models predict that performance across all 5 test formats (including comparisons between test formats with targets and novel foils as well as targets and similar lures; see Figure 11). In contrast, the CLS model predicts that there are many cases in which performance would not be correlated between test formats (Norman & O’Reilly, 2003). We observed clear evidence of relationships between performance on all of the test formats (see Figure 17), thus supporting the predictions of the distributed memory models. We note that Norman and O’Reilly (2003) also found that under conditions of high levels of encoding variability that the combined CLS model did predict there would be correlations between performance on different test formats. Thus, our finding of correlated performance between all 5 test formats is not necessarily discordant with all of the predictions the CLS model, but our results are naturally accounted for by the distributed memory models. It will be interesting for future studies to try to constrain encoding variability to test the strong prediction of the CLS model that the relationship between test formats would disappear in the absence of encoding variability; however, future computational modeling work should also examine how changes in encoding variability would change the predictions of the distributed memory models (cf. Huffman & Stark, 2017). Therefore, we hope that our empirical and modeling results will inform future research that seeks to explore the underlying mechanisms that explain the correlated performance that we observed here. We also think that our results provide an important cautionary tale about making inferences that these test formats (e.g., the comparison between test formats with similar lures compared to traditional recognition memory paradigms with novel foils; the comparison between the forced-choice test formats such as the A-A’ and A-B’ test formats) tap into fundamentally different cognitive or neural processes.

### Implications for Existing Models

Our results suggest that human memory is driven primarily by distributed representations (i.e., a better fit of the distributed memory models over the CLS hippocampal model); moreover, we previously found other results that support the distributed memory models. For example, we found that distributed memory models (MINERVA 2 and TODAM) could account for performance in both younger and healthy older adults on the A-X, A-A’, and A-B’ test formats (Huffman & Stark, 2017). Specifically, we found that the effects of healthy aging could be approximated by lowering the learning rate in the distributed memory models. Moreover, we recently showed that we could also account for memory differences on the A-X, A-A’, and A-B’ test formats in 4-year-olds, 6-year-olds, and young adults using an updated version of MINERVA 2 in which we fit the model to each participant’s data by allowing both the learning rate and encoding variability to change between each participant (Rollins, Huffman, Walters, & Bennett, 2023). We found that the best-fitting encoding-variability for both groups of children was significantly greater compared to the young adults. Moreover, we found that the best-fitting learning-rate for the 4-year-olds was significantly lower than for the 6-year-olds and the young adults. Importantly, we observed equivalent model fits for 4-year-olds, 6-year-olds, and young adults, thus suggesting that the differences in the best-fitting learning rates and encoding variability could not be explained by a confound of difference in overall model fit between groups. Additionally, we previously demonstrated that distributed memory models can account for the effects of repetitions on performance in the Mnemonic Similarity Task (Huffman & Stark, 2017). For example, previous research showed that young adults are more likely to false alarm to similar lures that were presented three times vs. once (Reagh & Yassa, 2014b), which led the authors to conclude that memory paradoxically degrades with repetitions. However, a signal-detection theory analysis of old/new with confidence ratings for targets and similar lures as well as the A-A’ test format revealed that performance was better for items that were presented three times vs. once (Loiotile & Courtney, 2015). Importantly, MINERVA 2 can account for both of these findings: repetitions increase the global match of the similar lure (thus accounting for the increase in false alarms in Reagh & Yassa, 2014b; Loiotile & Courtney, 2015) but the overall distributions become less overlapping (thus accounting for the improved performance from the signal-detection theory analysis as well as the A-A’ test format in Loiotile & Courtney, 2015; for more details see: Huffman & Stark, 2017). Altogether, our work suggests that the distributed memory models can account for a broad range of findings on the Mnemonic Similarity Task across multiple task conditions and provides mechanistic insight into differences in memory performance across the lifespan.

Our results provide a challenge for thinking about the role of individual regions in performing complex cognitive functions. Specifically, previous research with the CLS model has investigated simulated memory performance of different subregions of the brain, including the hippocampus and neocortex (O’Reilly & Norman, 2002; Norman & O’Reilly, 2003). If performance followed the pattern-separated representations of the hippocampus (and the dentate gyrus in particular), then we would expect a highly nonlinear transfer function between mnemonic discrimination and target-lure similarity in the input regions to the hippocampus. In contrast, if performance followed the more distributed representations of the neocortex, then we would expect a more continuous effect of target-lure similarity on mnemonic discrimination performance. Therefore, a more general question arises: How does the brain know which region to “listen” to when making memory decisions? Broader-scale neural network modeling could be implemented to allow us to get a better idea of what might happen within a more integrated neural network (e.g., that includes the prefrontal cortex, temporal lobe cortex, and hippocampus; cf. Norman & O’Reilly, 2003; Norman, 2010). For example, network-based theories of cognition suggest that cognitive performance is the result of interacting networks of brain areas and we cannot reduce cognitive functions to the level of single brain areas. For example, hippocampal damage in human patients was found to cause network-level disruptions (Henson et al., 2016). Moreover, studies of rodents have revealed that local lesions to the hippocampus cause widespread network changes (e.g., Jenkins, Amin, Brown, & Aggleton, 2006; Albasser, Poirier, Warburton, & Aggleton, 2007). Therefore, even lesion-based methods (arguably one of the methods most focused on studying localization of function) suggest that the hippocampus plays a network-based role in memory (we previously discussed these issues at length: Ekstrom, Huffman, & Starrett, 2017; Ekstrom, Harootonian, & Huffman, 2020). Distributed memory models provide a simplified framework that could offer fundamental insight into the nature of memory-based decision making: perhaps we combine or integrate all of the available information across the brain into a coherent memory trace (note that our ideas here also mesh well with neural theories of declarative memory: e.g., Squire et al., 2007; Wixted, 2007; cf. Norman, 2010).

The original goal of the CLS model was to provide a computational account for human memory performance on tasks that induce interference. For example, classic research showed that learning two overlapping lists of items within the AB-AC task caused some retroactive interference; however, participants tended to remember approximately half of the original AB list after learning the AC list (Barnes & Underwood, 1959). In contrast, feedforward neural networks using the delta rule almost completely forgot the AB list upon learning the AC list, an effect termed “catastrophic interference” (McCloskey & Cohen, 1989). More generally, catastrophic interference occurs when multiple lists of items are learned within neural networks using the delta rule (Robins, 2004). The original CLS model suggested that the finding of catastrophic interference, rather than being a downfall of this class of models, provided invaluable insight into the nature of how the brain might have evolved multiple memory systems. Specifically, McClelland et al. (1995) argued that the seemingly incompatible learning goals of episodic and semantic memory could reveal that the brain evolved separate learning systems to support each memory system: a fast-learning hippocampal system that employed pattern separation and a slower learning neocortical system that employed distributed representations (and delta-rule learning).

While the dual-process nature of the CLS model allows it to naturally account for differences in representations in the hippocampus vs. the neocortex, there are other computational frameworks that can also account for human performance on the AB-AC list learning task (as well as tasks that benefit from distributed representations). First, we note that the distributed memory models that we used here can account for human-like performance on the AB-AC task. For example, the Hopfield/autoassociative neural network (with Hebbian learning; cf. Robins, 2004) and TODAM (cf. Murdock, 1983; French, 1999) do not exhibit catastrophic interference. However, we readily acknowledge that the Hopfield/autoassociative neural networks are subject to other shortcomings, such as overall memory capacity constraints (e.g., Hopfield, 1982; note that newer versions overcome capacity limitations: Krotov & Hopfield, 2016). Notably, the storage capacity of Hopfield/autoassociative neural networks can be increased via delta-rule learning (Gardner, Wallace, & Stroud, 1989), but then the models exhibit catastrophic interference (e.g., Robins, 2004; but see methods for avoiding such interference below). Second, there are other computational frameworks that eliminate catastrophic interference within feedforward networks with the delta rule. For example, if these models are trained with interleaved learning, then they will not exhibit catastrophic interference (e.g., Robins, 2004; French, 1999; Robins, 1995; note that this was one of the central tenets of the CLS model: McClelland et al., 1995). Rehearsal is one method of interleaved learning in which the model is retrained on the original information during the learning of subsequent lists (e.g., French, 1999; Robins, 1995; Ratcliff, 1990). Although initial research in the 1950s provided evidence against a rehearsal-based framework for avoiding interference on the AB-AC list learning task (e.g., Barnes & Underwood, 1959), more recent research has suggested that rehearsal plays a critical role in reducing between-list interference. For example, behavioral research found that asking participants to retrieve the AB pairs during the AC list results in decreased proactive and retroactive interference (e.g., Hintzman, 2004; Wahlheim & Jacoby, 2013; Jacoby, Wahlheim, & Yonelinas, 2013; for review see: Kliegl & Bäauml, 2021). Moreover, fMRI studies have shown evidence that participants reactivate the original AB list during learning of the AC list (e.g., Zeithamova, Dominick, & Preston, 2012; Koen & Rugg, 2016; Schlichting, Guarino, Roome, & Preston, 2021), thus suggesting that the rehearsal of the AB list during the AC list could protect the memories from interference by allowing integration of the separate memories into a coherent memory trace. While there is behavioral and neural evidence supporting the idea that rehearsal strategies could help avoid catastrophic interference, there are other, similar methods that make weaker assumptions about the participants’ access to previously learned information. For example, pseudorehearsal is an alternate mechanism that could be used to avoid catastrophic interference and it consists of generating random input patterns and having the model learn whatever output is produced from its input (e.g., Robins, 2004, 1995). Thus, we could extend this framework to think of the learning phase during the AC list as having the participant simultaneously encode both whatever comes to mind when they see the trial stimuli, which could include at least a rough memory trace from the AB list (e.g., a form of recall-based encoding), as well as the actual presented stimuli. In summary, there are various computational frameworks that can account for performance on the AB-AC list learning task and we think it will be fruitful for future behavioral, neuroscientific, and modeling work to further explore these ideas and the connection between list-learning paradigms and tasks with targets and similar lures.

### Avenues for Testing the Generality of our Empirical and Modeling Results

While our results clearly suggest that human memory performance on the tasks with targets, similar lures, and unrelated foils is primarily driven by distributed representations, we think there are cases in which pattern-separated representations in the hippocampus might play a stronger role. For example, it is possible that task designs with so many trials could overload the pattern separation capabilities of the hippocampus. Participants in these types of paradigms view more than 100 images and have to make fine-grained decisions about them within a relatively short period of time (i.e., on the order of minutes to hours). Therefore, it is possible that under these circumstances, participants rely on the more distributed representations of the neocortex and that even the hippocampus exhibits a relatively high degree of pattern overlap (cf. Norman & O’Reilly, 2003; Norman, 2010). Future studies can vary the number of trials to see if performance varies as a function of list length, which would allow us to test the possibility that hippocampal pattern separation is overwhelmed by long lists (e.g., the transfer function between mnemonic discrimination performance and target-lure similarity in input regions of the hippocampus might be nonlinear with shorter lists [i.e., as opposed to our results that support a linear transfer function], the test-format effects might fit the CLS model’s predictions better for shorter lists, and the correlations between performance on the various tasks might diminish with shorter lists).

More recent theories suggest that event boundaries and schema knowledge exert a profound influence on human memory (e.g., Brunec, Moscovitch, & Barense, 2018). For example, the hippocampus appears to respond most prominently at event boundaries, thus suggesting that the hippocampus plays a key role in encoding memories around event boundaries (e.g., Baldassano et al., 2017; Reagh, Delarazan, Garber, & Ranganath, 2020; Reagh & Ranganath, 2023). Furthermore, a recent study found that event boundaries can influence mnemonic discrimination performance (Morse, Karagoz, & Reagh, 2023). Therefore, it is possible that memory paradigms that do not incorporate event-like structure might be harder for participants to parse, learn, and remember (i.e., because the sequence of images is essentially random and thus lacks event-like structure). It will be interesting for future experiments to combine more naturalistic approaches with the study of the transfer function between mnemonic discrimination and event similarity.

Another intriguing idea is that expertise can help us avoid interference between similar stimuli or memories. For example, people with domain expertise can rapidly detect and report differences between similar species of animals, similar faces, and other similar task stimuli such as Greebles (e.g., Tanaka & Curran, 2001; Gauthier & Tarr, 1997; Gauthier, Williams, Tarr, & Tanaka, 1998; Gauthier, Tarr, Anderson, Skudlarski, & Gore, 1999; Gauthier & Logothetis, 2000; Gauthier, Skudlarski, Gore, & Anderson, 2000; Mack, Love, & Preston, 2016). Likewise, in our daily lives, we might be experts in our knowledge of typically recurring experiences (e.g., between events that contain similar, familiar people and places). Thus, it is possible that schema knowledge could aid in our ability to distinguish between similar experiences. Therefore, it will be interesting for future research to examine whether more naturalistic designs, that incorporate event schemas and expertise, confer protection from interference between similar memories as well as the neural mechanisms that support behavioral performance under these conditions.

A related possibility is that the hippocampus might play a stronger role in the pattern separation of memories that are inherently associative, such as spatial or temporal memories. Several prominent theories suggest that the hippocampus plays a key role in binding items in context or in spatial memory more generally (e.g., Eichenbaum et al., 2007; Ranganath & Ritchey, 2012). For example, previous research has provided evidence of pattern separation between similar spatial environments in rodents (J. K. Leutgeb et al., 2007; Neunuebel & Knierim, 2014) and humans (Kyle, Stokes, Lieberman, Hassan, & Ekstrom, 2015; Stokes, Kyle, & Ekstrom, 2015). Likewise, future research could further leverage temporal memory paradigms (e.g., Roberts, Ly, Murray, & Yassa, 2014; Reagh et al., 2016; Montchal, Reagh, & Yassa, 2019) to investigate whether there is a more linear relationship between mnemonic discrimination and hippocampal activity patterns. Therefore, it is possible that people would exhibit less interference for distinguishing between similar stimulus-stimulus associations, spatial memories, or temporal sequence memories. Future research could leverage novel advances in virtual reality technology to address these possibilities, especially regarding changes in spatial environments, which could be based on previous neuroscience research (J. K. Leutgeb et al., 2007; Neunuebel & Knierim, 2014; Kyle et al., 2015; Stokes et al., 2015).

## Conclusion

We generated and tested predictions of distributed memory models vs. the standard CLS hippocampal model on several test formats that included targets, similar lures, and novel foils. All of our empirical data were better fit by the predictions of the distributed memory models, thus suggesting that memory performance is primarily driven by distributed representations (e.g., neocortex, CA1, CA3) rather than the pattern-separated representations of the dentate gyrus or a composite representation across these systems. Moreover, our results provide important insight into influential theories about the cognitive neuroscience of memory; for example, our results provide new insight into longstanding debates in the field regarding the contributions of the hippocampus vs. neocortex to human memory performance. Thus, our behavioral and computational modeling results provide novel avenues for future behavioral, neuroscientific, and computational modeling research.

## Author Contributions

Conceptualization: DJH. Data Curation: RG and DJH. Formal Analysis: DJH and RG. Funding acquisition: DJH. Investigation: RG and DJH. Methodology: DJH and RG. Project administration: DJH. Resources: DJH and RG. Software: DJH and RG. Supervision: DJH. Visualization: DJH and RG. Writing - Original Draft: DJH. Writing - Review & Editing: DJH and RG.

## Notes

We have no competing interests to disclose.

This work was supported by startup funds from Colby College as well as funds from the Department of Psychology. A small part of the initial work with the computational models was partially supported by National Institute of Mental Health grant F32MH116577 to DJH.

### Competing Interest Statement

The authors have declared no competing interest.

### Summary of Updates

We made updates to a few key aspects of our paper: 1) we updated the framing of the paper to place a greater emphasis on the importance of our findings, 2) we added new analyses to allow us to more clearly test the predictions of various models, 3) we updated the format of the paper to make our paper more compartmentalized and hopefully clearer.

